# Attention network for predicting T cell receptor-peptide binding can associate attention with interpretable protein structural properties

**DOI:** 10.1101/2023.02.16.528799

**Authors:** Kyohei Koyama, Kosuke Hashimoto, Chioko Nagao, Kenji Mizuguchi

## Abstract

Understanding how a T cell receptor (TCR) recognizes its specific ligand peptide is crucial for gaining insight into biological functions and disease mechanisms. Despite its importance, experimentally determining TCR-peptide interactions is expensive and time-consuming. To address this challenge, computational methods have been proposed, but they are typically evaluated by internal retrospective validation only, and few have incorporated and tested an attention layer from language models into structural information.

Therefore, in this study, we developed a machine learning model based on a modified version of the Transformer, a source-target-attention neural network, to predict TCR-peptide binding solely from the amino acid sequences of the TCR’s complementarity-determining region (CDR) 3 and the peptide. This model achieved competitive performance on a benchmark dataset of TCR-peptide binding, as well as on a truly new external dataset. Additionally, by analyzing the results of binding predictions, we associated the neural network weights with protein structural properties. By classifying the residues into large and small attention groups, we identified statistically significant properties associated with the largely attended residues, such as hydrogen bonds within the CDR3. The dataset that we have created and our model’s ability to provide an interpretable prediction of TCR-peptide binding should increase our knowledge of molecular recognition and pave the way to designing new therapeutics.

## INTRODUCTION

The T cell receptor (TCR) serves as an antigen receptor, primarily composed of alpha (TCR*α*) and beta (TCR*β*) chains. It has a remarkable sequence diversity in its complementarity-determining regions (CDRs), similar to the B cell receptor, antibody. The TCR’s CDR3, found in both *α* and *β* chains (CDR3*α* and CDR3*β*, respectively), is the most diverse and vital for recognizing antigenic peptides presented by the major histocompatibility complex (MHC) molecule. Given the immense sequence diversity produced through somatic recombination, the TCR’s potential interactions with different peptides are enormous. Therefore, predicting TCR-peptide binding, primarily involving CDR3, is of great importance. This prediction could significantly impact our understanding of biological functions and disease mechanisms, and guide potential disease recovery pathways.

In response to this, there have been numerous machine learning methods developed for TCR-peptide prediction (Gowthaman and Pierce, 2019; Montemurro et al., 2021; Dash et al., 2017; Springer et al., 2020, 2021; Lu et al., 2021a; Gao et al., 2023). Some studies in the bioinformatics field were in line with models using the source-target attention (Chen et al., 2020; Weber et al., 2021; Koyama et al., 2020; Honda et al., 2020), and there are researches attempting to apply the attention models to the TCR-peptide binding prediction (Wu et al., 2021; Sidhom et al., 2021; Xu et al., 2022, 2021).

The Transformer (Vaswani et al., 2017) and BERT (Devlin et al., 2018) models, known for their impressive results and interpretability (Voita et al., 2019; Hao et al., 2021; Rogers et al., 2020), have demonstrated the advantages of the crossattention mechanism in source-target multi-input tasks such as machine translation or image-text classification (Gheini et al., 2021; Parthasarathy and Sundaram, 2021; Lee et al., 2018). Furthermore, during the training process, employing the crossattention mechanism on two separate sequences is less computationally intensive than applying a self-attention model to concatenated sequences. This is because the computational complexity of the Transformer’s attention mechanism scales quadratically with the length of the input sequence. Despite the wide application of the Transformer, a comprehensive analysis of interpretability based on the multi-input TCR-peptide protein complex is yet to be provided. Few studies have tried to provide the source-target-attention model of the Transformer at the level of individual residues in the CDR3*α β* or peptide and analyze structural information such as hydrogen bonds.

For instance, models such as NetTCR-2.0 (Montemurro et al., 2021) and ERGO-II (Springer et al., 2021), despite demonstrating impressive predictive capabilities, are based on convolutional or recurrent neural network frameworks. The PanPep model (Gao et al., 2023), while using an attention mechanism, focuses solely on the CDR3*β*. This model provides no information about the structurally important residues on the alpha chains, and it does not account for interaction factors related to hydrogen bonds. The TCR-BERT (Wu et al., 2021) model uses both the alpha and beta chains. However, it is trained without the peptides and does not map the attention on residues for structural analysis. The model proposed by AttnTAP (Xu et al., 2022) utilizes attention but it does not directly use the Transformer attention on both sides of TCR and peptide. It does not incorporate the alpha chain, either. DLpTCR (Xu et al., 2021), another model in this field, employs ResNet attention; however, it refrains from using the Transformer attention.

Unlike existing research, in essence, our model intends to develop a computational method that can incorporate CDR3*α*, CDR3*β*, and peptide, and conduct a residue-wise structural analysis, leveraging a Transformer-based attention mechanism on sequences. We hypothesize that an attention-based neural network can accurately predict TCR-peptide binding and provide interpretable biological insights into the TCR function. To achieve this purpose, we propose a model, the Cross-TCR-Interpreter, which uses a cross-attention mechanism for predicting TCR-peptide binding, the binding between CDR3 regions of both the *α* and *β* chains, and a peptide.

Our model achieved competitive performance on the benchmark. Also, by performing statistical tests on the attention values over the complex structures, we successfully identified statistically significant structural properties of largely attended residues such as hydrogen bonds and residue distance. We will also discuss the limitations of generalizability on unseen data, an issue not unique to our model but is also evident in other models. Our approach, leveraging the Transformer’s sourcetarget-attention neural network, highlighted the capacity for a deeper understanding and analysis of protein interactions.

## MATERIALS AND METHODS

### Model

An overview of the prediction model used in this study is shown in Figure 1. The peptide sequence and the sequence of CDR3*α* and CDR3*β* connected with the connection-token (colon “:”) were processed separately in the embedding layer and Transformer, and then they were input into the cross-attention layer designed for interaction prediction. The cross-attention was used to create a mutual-only layer, enabling the model to verify the interactions. The outputs of the cross-attention layer were concatenated and averaged over the length direction in the output layer. A multi-layer perceptron (MLP) layer outputs a single prediction as a real value, known as the confidence value, from 0 to 1, whereas a true binding datum is represented as a binary value of 0 or 1. Binary cross entropy (BCE) was used as the loss function, and the model output was evaluated using the ROCAUC score and the average precision score.

**Figure 1.**
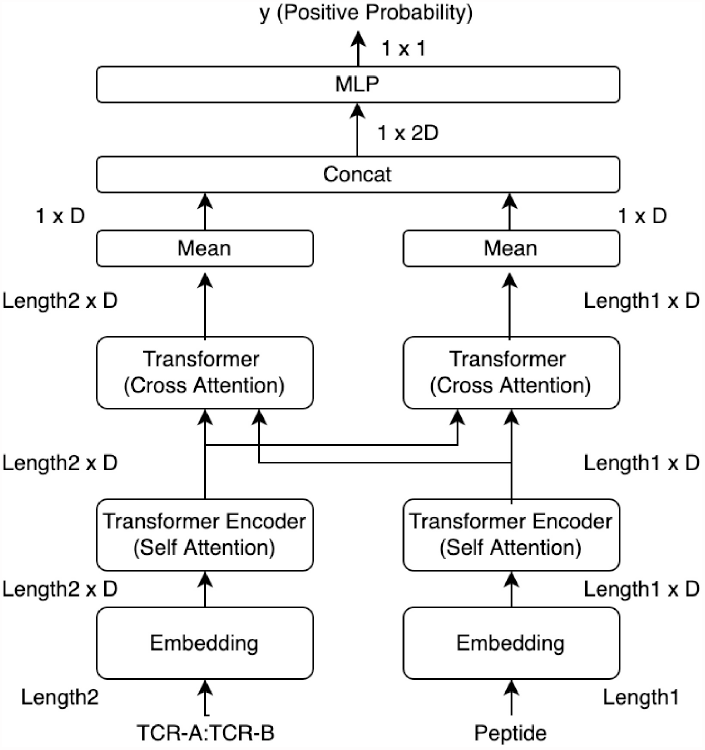
Overview of our Cross-TCR-Interpreter model. Data tensor sizes are denoted. The crossattention layers in the middle of the figure were analyzed using structural data after being trained with sequence data. Each embedding layer takes amino acid sequences as input.

The model only takes amino acid sequences of CDR3*α*, CDR3*β*, and peptide as inputs. We used only the CDR3s and not the entire TCR sequences. Any other information such as gene types is not utilized. Leveraging solely sequence information, without incorporating domain-specific human knowledge such as gene or MHC information, should be surely the key part for emulating interpretability, closely resembling the natural phenomena of CDR3 binding. The CDR3 and peptide sequences were represented by 20 amino acid residues. Positional embedding and padding tokens were also added to the sequences. Padding was performed so that the lengths of each CDR3 sequence aligned with the maximum sequence length in the training data. The same was performed for the peptide side. The CDR3*α* sequences had a maximum length of 28 and a minimum length of 7. Similarly, the CDR3*β* sequence length ranged from 5 to 28. For the peptides, the sequence length ranged from 8 to 25.

The cross-attention layer is a modified model of Transformer attention. In particular, it takes two sequences as input values and allows meaningful information to be extracted from the entire information of one sequence based on the entire information of the other sequence, implying that it is beneficial for interaction predictions.

The attention layer is specified by the following Equation 1.

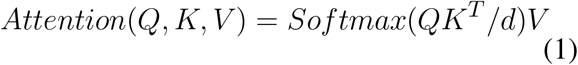

In Equation 1 for the attention layer, *𝒬, K*, and *V* are the data matrices of sequences, and *d* is the scaling factor. *K*^*T*^ denotes transposed matrix of *K*, where the sizes of arrays are *Q* : *L*_1_ *×D, K* : *L*_2_ *×D*, and *V* : *L*_2_ *×D. D* is the embedding dimension. When *Q* = *K* = *V* and *L*_1_ = *L*_2_, this is a self-attention layer.

In the cross-attention layer, *K*(= *V*) and *Q* represent two different inputs, i.e., a connected sequence of CDR3*α* :CDR3*β* and a peptide, respectively.

In addition, we defined four heads for each side in the cross-attention layer and those heads were concatenated as in a typical Transformer. The softmax function defines the weights to *V* when matrix *Q* is input, and the weights are allocated so that the sum is 1 over the length direction of *V*. This *Softmax*(*QK*^*T*^ */d*) is the attrition and is used for the analysis and visualization, suggesting the residue positions that are important within the length of *V*.

By representing the learned hidden values of the CDR3s, taken from the output of the cross-attention layer just before concatenation, as *H*_*T CR*_, we have:

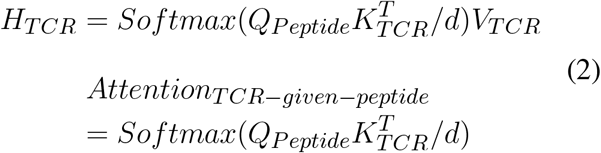

The same was done for the peptide side and we obtained 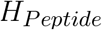. 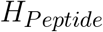 and *H*_*TCR*_ were concatenated just before the MLP layer.

Visualization and analysis of the attention layer for interactions allow interpretation of the interaction across sequences (e.g. Figure 4). The cross-attention layer uses peptides as inputs and assigns specific weights to each residue of CDR3s to learn the important sites of CDR3s, and vice versa. This enabled us to analyze each side of the two areas of attention separately.

All hyperparameters of the model are tuned with the Optuna package (Akiba et al., 2019) and reported in Supplementary Information. Except for the hyperparameter tuning, the training was completed with one A100 GPU node at the Osaka University SQUID cluster for approximately 3 hours, and the inference was completed with a 2.6 GHz 6-Core Intel Core i7 CPU for approximately 2 hours.

### Preparation of training and test datasets of sequences

For the CDR3s and peptide binding datasets, we took the repository of ERGO-II (Springer et al., 2021) which contains McPAS (Tickotsky et al., 2017) and VDJdb (Shugay et al., 2018). We also independently downloaded and created the newer version of VDJdb and Covid-19 datasets (Lu et al., 2021b). In more detail, the sequence datasets we have created are as follows:

- **Benchmark datasets McPAS and VDJdb-without10x (training and test):** The two primary benchmark datasets, McPAS and the VDJdb were derived from ERGO-II repository. Specifically, the VDJdb dataset excluded the 10x genomics data (referred to as VDJdb-without10x). These datasets had both training and test sets and contained both positive and negative interactions.
- **Entire data dataset (training):** For a more comprehensive model training, we also utilized an extended dataset, referred to as the “entire data”. This dataset concatenated the McPAS dataset, VDJdb-without10x, and VDJdb with the 10x genomics data (VDJdb-with10x). This dataset was used for training the model to evaluate the following recent data test set and the Covid-19 dataset.
- **Recent data test set (test):** To assess the entire-data-trained model’s effectiveness in handling new, unseen data, we tested our model on a recent test set from VDJdb downloaded in 2023. The negative interactions were added by randomly choosing the CDR3s and peptides, as the data contains only positive interactions. This evaluates whether a model performs on the most up-to-date data, highlighting its predictive capability for new TCR-peptide interactions.
- **Covid-19 dataset (test):** Lastly, to provide a stringent assessment of our entire-data-trained model, we created a dataset derived from the study on Covid-19 (Lu et al., 2021b). This dataset is recognized as one of the most challenging for models trained on the entire data dataset.

As this study involves a binary classification problem, negative label data were needed to train the model. However, as most of the TCR and peptide response data are positively labeled, this study followed the same configuration and data of the existing ERGO-II report that generated random TCR-peptide pairs and assigned negative labels to adjust the positive-to-negative ratio. The size of the negative data was five times larger than that of the positive data. Therefore, each data record to train the model is a tuple of CDR3*α*, CDR3*β*, peptide that has a binary label. When either the CDR3*α* or CDR3*β* sequence was missing for an interaction pair, the interaction pair was removed and not used for training.

To ensure no overlapping pairs, we meticulously eliminated the CDR3*α*, CDR3*β*, peptide pairs from the test set that were present in the training set. However, there may still exist duplicated pairs of CDR3*α*, CDR3*β* or individual CDR3 or peptides that appear in both training and test sets, because the same TCR is present in both the test and training data sets and may be paired with other different multiple peptides. The proportion of such duplicates for McPAS and VDJDB is described in the Result section.

### Benchmark dataset and experiment

The validity of the cross-attention model was confirmed by comparing the test scores on the benchmark data using McPAS and VDJdb without 10x Genomics data (VDJdb-without10x). The benchmark models included ERGO-II (Springer et al., 2021) and NetTCR2.0 (Montemurro et al., 2021), which use both CDR3*α* and CDR3*β*. The only CDR3*β* chain TCR-peptide prediction models such as NetTCR2.0(Montemurro et al., 2021), PanPep (Gao et al., 2023), AttnTAP (Xu et al., 2022), and DLpTCR (Xu et al., 2021) were also compared. In these data, the binary labels were assigned to CDR3*β* and peptide pairs. We evaluated our model performance not only by using the whole test set but also by using the per-peptide score within the test set. The benchmark datasets in the existing ERGO-II research were developed by incorporating assumed negatives, followed by splitting them into training and test datasets. This approach might create an oversimplified problem, as many peptides or CDRs are likely to be shared between the training and test datasets.

The detailed benchmark dataset creation process was as follows:

- Step 1: Download the test set and training set of ERGO-II and remove interactions that do not have either one of CDR3*α*, CDR3*β*, or peptide.
- Step 2: Remove duplicated interaction pairs from the test set that are shared with the training set.

While training, we minimized the binary cross entropy for the training set in the benchmark experiments. If the binary cross entropy did not improve within 10 updates, we stopped the training. Subsequently, we adopted the weights that gave the minimum value of the binary cross entropy as the best model.

### The entire data dataset and the recent data test set

After confirmation of the model’s performance, we trained the model again with the whole dataset (namely, the “entire data” dataset) that included McPAS, VDJdb-without10x, and VDJdb-with10x. Our primary objective with the entire data dataset approach was to uncover meaningful relationships and model the binding nature of the interactions, potentially leading to a meaningful interpretation. By using this entire-data-trained model, we expected to acquire the learned relationship between the two sequences within the attention layer. The entire data dataset included 10x dataset (10x Genomics, 2019) that was omitted in the benchmark experiments. By using all the data, we attempted to incorporate the maximum possible information related to binding into the model, and herein to analyze the attention weights in the trained model. For the purpose of this model, we designated the test set to comprise the most recent data from VDJdb (namely, the “recent data” test set), specifically the data downloaded between 2022 and 2023. In contrast, the training set included data downloaded from VDJdb prior to 2022 and McPAS data. After downloading the data, we added five times more negative interactions to the downloaded recent data test set. These negative pairs of CDR3s and peptides are sampled only from the recent data test set, not from the entire data dataset. This recent data test set can resemble a realistic situation where we use the model with prospective validation, evaluating the model non-retrospectively.

The detailed entire data dataset and the recent data test set creation processes are as follows:

- Step 1: Download the McPAS, VDJdb-without10x, and VDJdb-with10x data of ERGO-II. Concatenation and removal of interactions that do not have either one of CDR3*α*, CDR3*β*, or peptide.
- Step 2: Remove duplicated pairs inside the dataset (the entire data dataset).
- Step 3: Download the VDJdb in June 2023 and create the pairs of CDR3*α*, CDR3*β*, and peptide (the recent data test set).
- Step 4: Remove duplicated interaction pairs from the recent data test set that are shared with the training set.
- Step 5: Add five times more negative interactions to the recent data test set.

The difference between the recent test and benchmark datasets lies in the timing of the data split. For the recent data test set, we performed the data split prior to adding assumed negative samples to avoid the issue of the oversimplified problem. To show how diverse the recent data and the entire data dataset were, the sequence-sequence pairwise distance matrix was calculated using Clustal Omega software (Sievers et al., 2011) for sequence space analysis.

### Covid-19 datasets and experiment

Similarly to the recent data test set, to evaluate how accurately the entire-data-trained model would perform in a realistic situation that has no known peptides, we applied it to prediction tasks of a real-world Covid-19 dataset generated from the Covid-19 study (Lu et al., 2021b). A virtual dataset was created using the TCRs pairs and peptides taken from the S(Spike) protein. In the original study, the reaction between the peptides and TCRs was evaluated by a reporter cell assay by measuring green fluorescent protein expression in the pathway of TCR, and the peptides of the S protein were created with a 15-length residue window of amino acid residues by moving four strides of residues. We adopted the same procedure virtually to create the peptides of the S protein, by creating a 10-length residue window and moving one residue stride, as the median length of peptides in the training dataset was 10. There was no peptide overlap between the entire data dataset peptides and the 10-length peptides of the Covid-19 dataset. To demonstrate the diversity of the Covid-19 peptides, we computed the sequence-sequence pairwise distance matrix in the same manner as we did for the entire data dataset.

### Attended residue analysis with attention values on 3D structures

After training the model on the entire data dataset, we could acquire any attention matrix on arbitrary residues. We argue that it makes sense to analyze the model since we used the data that are correctly predicted. Our approach was not aimed at cherry-picking but rather at investigating and interpreting significant features discerned by the model.

Thus, our goal was akin to discerning the impact of age on a mathematics test score through a least squares regression analysis. In such an analysis, our principal interest would lie in the correlation and its magnitude. Therefore, we could determine that age has a correlation with the math score, while sex or nationality do not. Similarly, in our research, the primary focus was to comprehend the crucial features, or attention values, that our model has learned.

Dividing the residues into two groups of large and small attention made it possible to analyze the attention values. For each head of CDR3s attention provided a peptide, we defined the residue indices of large CDR3s as *R*_*large,h*_ in Equation 3.

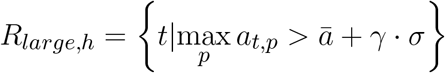

where *h* denotes head and *a*_*t,p*_ denotes an attention value of CDR3 residue index *t* and peptide residue index *p*.

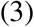

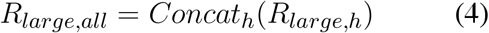

Equation 3 shows the TCR side attention. Given a head *h* of the cross-attention layer, let *A*_*h*_ be an attention matrix of the TCR side with elements *a*_*t,p*_. Please note that, by the definition in Equation 1, Σ_*t*_ *a*_*t,p*_ is one dimensional all-one vector, (1_1_, 1_2_,, 1_*p*_,, 1_*P*_), where *P* is the peptide’s length. We defined this as TCR-side attention because each *p* assigns the attention to TCRs as a sum of one. max_*p*_ takes the maximum value to the peptide axis. *ā* is the mean of the attention values of *A*_*h*_, and *σ* is the standard deviation of *A*_*h*_. *γ* is a factor that defines the large or small definition that is empirically expressed later in the Result section. When computing the largely attended peptide residues, we exchanged the notation of *t* and *p*. When computing the not-largely attended residues, we replaced the in-equation operator with smaller-than. Equation 4 shows the attended residues of the TCR side when all heads are concatenated.

### Protein Data Bank (PDB) structure data analysis

Defining the largely attended residues, the results were examined using a dataset of TCR-peptide complex structures taken from the Protein Data Bank (PDB) (Berman et al., 2003). We collected TCR-related structures from PDB Search and the SCEptRe server (Mahajan et al., 2019), which gathers the complex structures of TCRs. The SCEptRe data used here in this study were downloaded on June 2, 2021. With PDB headers, 65 structures with alpha and beta chains were identified. Anarci (Dunbar and Deane, 2016) was used to extract the CDR3 portion of the structures. These 65 were narrowed down to 55 by setting restrictions on the lengths of the TCRs and peptide sequences. The 55 structures contained eight pairs with identical sequences for the CDR3s and peptides, and therefore, a final analysis was performed based on the sequences of 47 structures.

We performed a paired student’s t-test (also called the dependent t-test) to assess the differences between the two residue groups of largely attended and not-largely attended. The paired t-test is a statistical method used to compare the means of the two groups of subjects that are dependent on each other. In this study, the TCR-peptide complex structures were used as subjects of the t-test. The test values of the t-test were properties such as the proportion of TCR residues that were hydrogen-bonded to the peptide, whether the residue was engaged in an H-bond, or how many H-bonds the residue had. We used BioPython (Chapman and Chang, 2000) and LIGPLOT (Wallace et al., 1995) to gather the structural properties.

### Input Perturbation

To examine individual cases in greater detail, we employed the input perturbation method, which evaluates the sensitivity of a model to changes in its inputs. This approach complements the broader understanding provided by the paired t-test on the group.

The input perturbation method involves substituting amino acid residues at some critical positions with alternative amino acids and observing the resulting changes in both prediction and attention values. By altering the attended residues, we assessed the model’s responsiveness to these modifications, offering the observation of the changes in predictions and attention values.

## RESULTS

### Study overview and experiment types

We performed three experiments to validate the performance and usefulness of our proposed Cross-TCR-Interpreter model (Figure 1). In the first experiment, we trained and validated the model using existing benchmark datasets, comparing its performances with those of previously proposed models. In the second experiment, in order to conduct the external prospective validation of TCR-peptide binding, we retrained the model using the entire data dataset and validated it with the Covid-19 dataset (Lu et al., 2021b) and the recent data test set. In the third experiment for explainability, we applied the entire-data-trained model to a dataset of TCR-peptide pairs of known 3D structures, performing statistical analyses of cross-attention values to detail the biochemical binding event. Also, we have used the model for the input perturbation analysis to observe the change of attention. Hence, although the model was exclusively trained on sequence data, the interpretation of its predictive modeling was further enhanced using structural data.

### Unique element overlap and interaction-wise overlap can explain the difficulties of datasets

Our sequence datasets’ key statistics are summarized in Table 1. The training interactions of benchmark datasets are 23,363 for McPAS and 19,526 for VDJdb-without10x. The interactions of test sets are 4,729 for McPAS and 4,010 for VDJdb-without10x, with no duplicates between the test and the training data.

**Table 1.** Dataset statistics. The “Interaction” column means the unique count of pairs of *{*CDR3*α*, CDR3*β*, Peptide*}* and CDR3*α β* denotes the unique count of pairs of *{*CDR3*α*, CDR3*β}*. The duplication count, the “in duplication” row of the “Unique count” column, means the number of unique data that are shared between training and test sets, i.e. overlapped data count. The “Pos. Rate” column denotes the positive ratio in the interaction.

| Dataset name | Unique count | $CDR3\alpha\beta$ | Peptide | Interaction | Pos. Rate |
| --- | --- | --- | --- | --- | --- |
| McPAS | in training | 3181 | 316 | 23363 | 0.1665 |
| McPAS | in test | 833 | 190 | 4729 | 0.1512 |
| - | in duplication b/w training and test | 132 | 171 | 0 | N/A |
| VDJdb-without10x | in training | 2902 | 175 | 19526 | 0.1670 |
| VDJdb-without10x | in test | 689 | 120 | 4010 | 0.1504 |
| - | in duplication b/w training and test | 111 | 111 | 0 | N/A |
| The entire data dataset (A) | in training | 23299 | 478 | 119046 | 0.1400 |
| The recent data test set (B) | in test | 33183 | 838 | 33360 | 0.1667 |
| The Covid-19 dataset (C) | in test | 1676 | 1264 | 2118464 | $1.4 \cdot 10^{-5}$ |
| - | in duplication b/w (A) and (B) | 18 | 44 | 0 | N/A |
| - | in duplication b/w (A) and (C) | 1 | 0 | 0 | N/A |

In the table, unique counts of CDR3s, peptides, and interactions are displayed. Specifically, for instance, the McPAS training set consists of 23,363 interactions involving 3,181 unique CDR3s sequences and 316 unique peptides, with 16.67% being positive. From these unique sequences, 833 CDR3s and 190 peptides also appear in the test dataset, whereas none of the same interaction pairs of CDR3s-peptide appear there in the test set. Under ideal circumstances, full observations between these unique CDR3s sequences and unique peptides would have yielded an interaction count of 1,005,196 (= 3181*·*316). However, due to data limitations in real-world datasets, this situation is not realized. There are very few overlapped duplications on CDR3s and peptides between the entire data dataset and the recent data test set. Also, there are very few duplications of CDR3s and peptides between the entire data dataset and the Covid-19 dataset. This explains the difficulty in predicting the interactions in the recent data test set and the Covid-19 dataset.

Figure 2 outlines interaction-wise duplication within each test dataset, meaning the duplication count of interaction where one side of peptide or CDRs is shared with the training dataset. As shown, the test set interactions of the McPAS and VDJdb-without10x are composed of already seen peptides in the training dataset, while the 14.8% interactions of the recent data test set comprise known peptides and no interactions of the Covid-19 dataset peptides are seen in the entire data dataset. From both perspectives of peptides and CDRs, the recent data test set and the Covid-19 dataset show interactions mostly of unseen CDRs or peptides.

**Figure 2.**
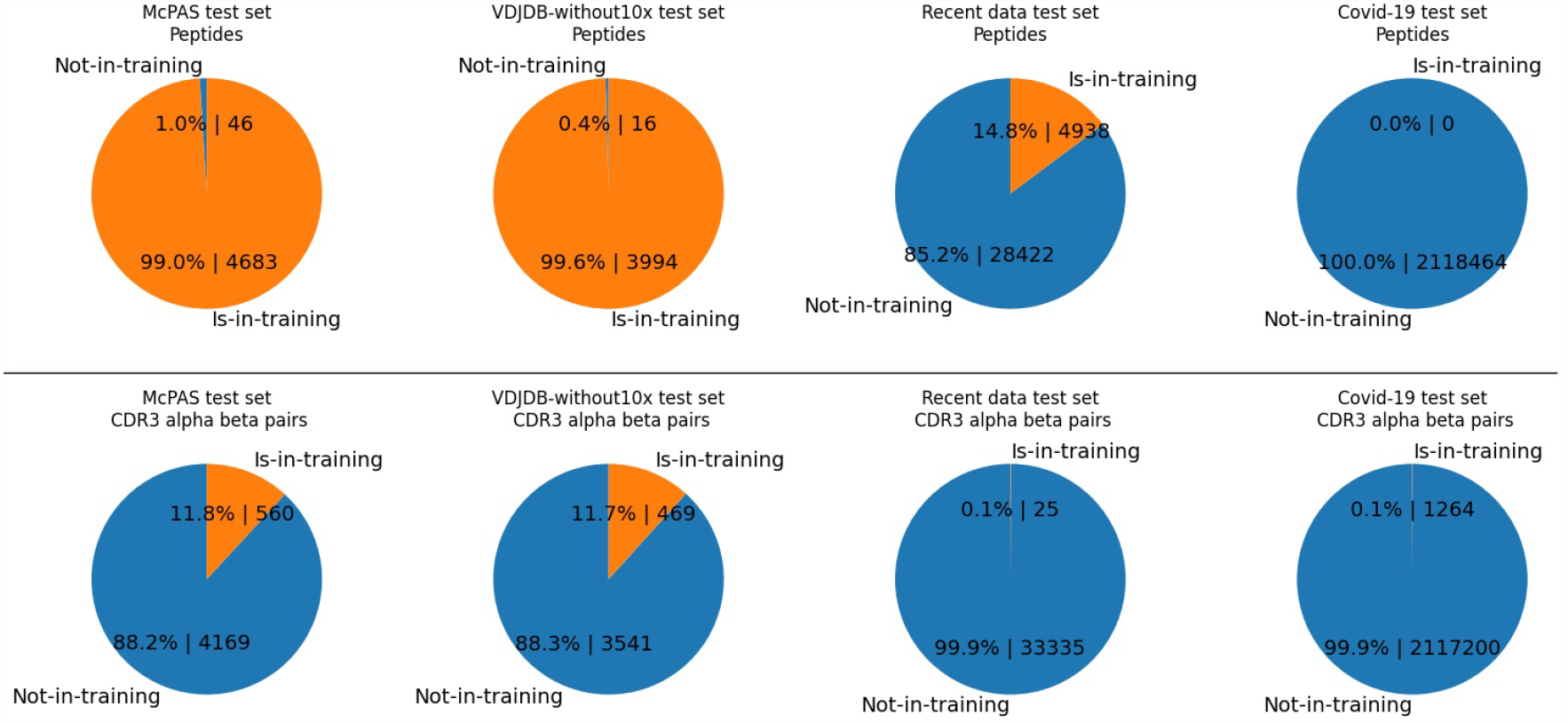
The pie charts illustrate percentages and counts of interactions of the test set comprising either one of seen elements. Contrary to the Table 1, the numbers are the duplication counts of interaction where one side of peptide or CDRs is shared with the training dataset In each test set, we display either the duplicated counts of CDR3 pairs or the duplicated counts of peptides. ‘Is-in-training’ indicates the peptide or CDR3s are present in the training dataset, while ‘Not-in-training’ means the peptide or CDR3s are not found in the training dataset. The test set interaction counts of McPAS is 4,729, that of VDJdb-without10x is 4,010, that of the recent data test is 33,360, and that of the Covid-19 test set is 2,118,464. Upper half row: Peptides. Lower half row: CDR3*α β*. Each column shows a different data set; from the left, they are McPAS, VDJdb-without10x test set, the recent data test set, and the Covid-19 test set. For example, the McPAS test set consists of 4,729 interactions, in which 4,683 interactions are comprising peptides seen in the training set and 46 interactions are comprising brand new peptides. From the CDR3s aspect, 560 interactions out of 4,729 are composed of unseen CDR3s, whereas 560 interactions are composed of seen CDRs.

For instance, the McPAS test set consists of 4,729 interactions, of which 4,683 interactions are comprising peptides already seen in the training set and 46 interactions are comprising brand new peptides. However, the recent data test set consists of 33,360 interactions and only 4,938 interactions comprise the peptides seen in the training set.

### The model shows excellent performance for benchmark datasets

To evaluate the performance of our model, we used training and test datasets inspired by those of ERGO-II (Springer et al., 2021). Two benchmark datasets, McPAS and VDJdb without 10x Genomics data (VDJdb-without10x), were prepared for this experiment. Evaluating the models with the ROCAUC score and the average precision score, our model showed competitive scores against other models for both benchmark datasets in the sequence-feature-only setting models (Table 2 and Table 3).

**Table 2.** Result of benchmark dataset of McPAS. APS stands for Average Precision Score.

| Model | Features in addition to peptides | ROCAUC | APS |
| --- | --- | --- | --- |
| Our model | CDR3s of $\alpha$ and $\beta$ chains | 0.9154 | 0.6211 |
| NetTCR2.0 | CDR3s of $\alpha$ and $\beta$ chains | 0.9204 | 0.5808 |
| PanPep | CDR3 Sequence of $\beta$ chain with biochemical features | 0.8374 | 0.4519 |
| AttnTAP<sup>1</sup> | CDR3 Sequence of $\beta$ chain | 0.840 | - |
| DLpTCR<sup>1</sup> | CDR3 Sequence of $\beta$ chain | 0.633 | - |
| ERGO-II, LSTM<sup>2</sup> | CDR3s of $\alpha$ and $\beta$ chains | 0.855 | - |
| ERGO-II, LSTM<sup>2</sup> | CDR3s of $\alpha$ and $\beta$ chains, VJ genes and MHC Type | 0.939 | - |

**Table 3.**
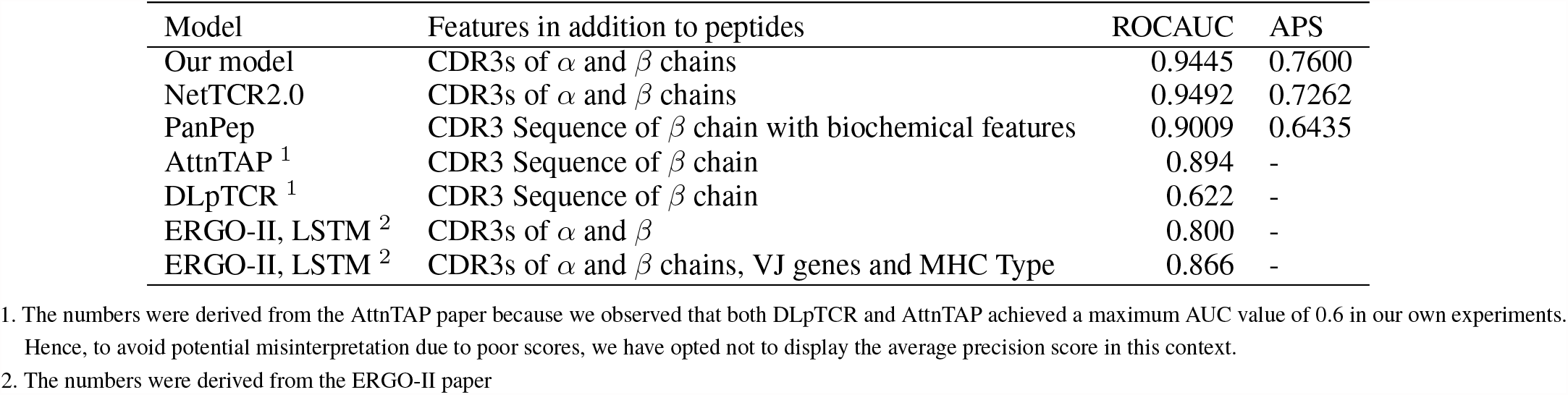
Result of benchmark dataset of VDJdb-without10x. APS stands for Average Precision Score.

For detailed performance metrics per-peptide for each test set, we have calculated the scores on the top eight frequent peptides in Figure 3. Our model shows competitive results over the NetTCR2.0(Montemurro et al., 2021) model for the per-peptide performance comparison. We added an analysis of the performance delegation of TCR distance in the Discussion section.

**Figure 3.**
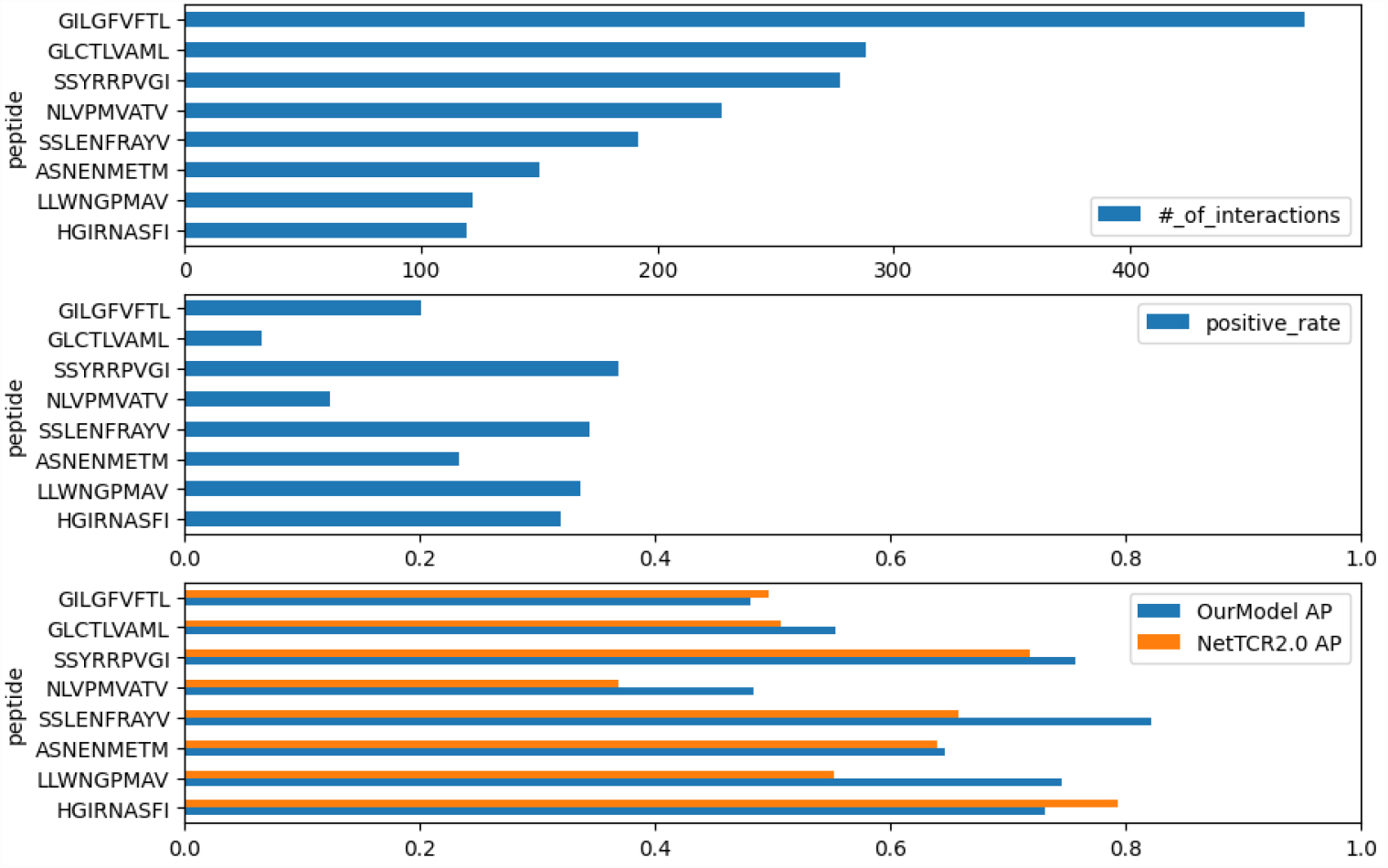
The number of interactions for each peptide, the positive rate inside the interaction (the ratio of positively recorded CDR3*α β*), and the average precision scores (AP) in the benchmark data of McPAS.

**Figure 4.**
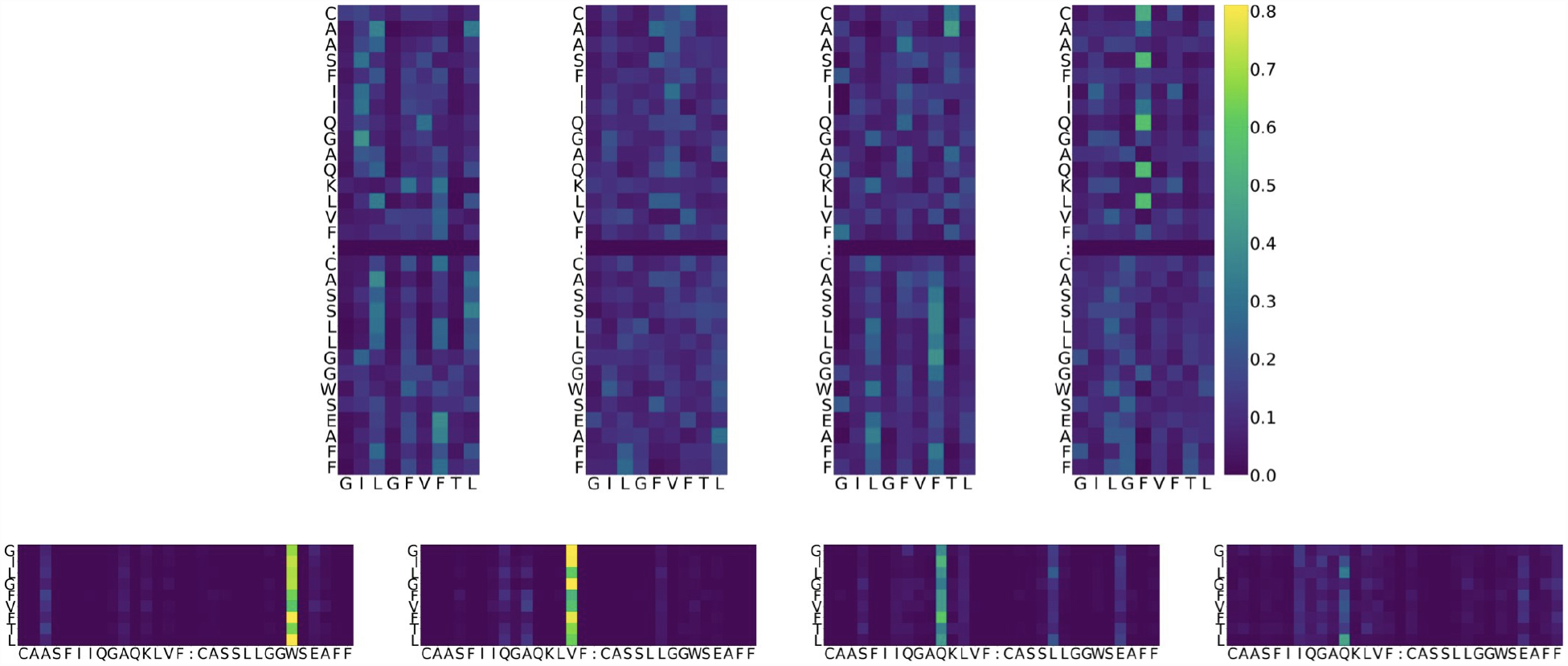
Example of attention value visualization of PDB ID 5TEZ (Yang et al., 2017). Upper half: attention values of a peptide so that the sum is 1 over the peptide given a CDR3*α β* pair. The X-axis represents the residue of the peptide, while Y-axis represents the residue of TCRs. Lower half: attention values of a CDR3 pair so that the sum is 1 over the CDR3s given a peptide. The X-axis is the residue of TCRs while Y-axis is the residue of peptides. The sum over the x-axis direction is 1 for both images. Four columns denote the heads of the multi-head attention layer. Colors denote the magnitude of attention value: dark blue represents smaller attention, yellow represents larger, and green is in the middle. In the lower figure, the cell corresponding to peptide position *L*_8_ (the last row) and CDR3*β* position *W*_24_ (sixth column from the right) represents the weight of how important the CDR3 *W*_24_ is given the peptide *L*_8_. Its color is bright yellow, which means that the attention value is large and the two residues might play a potentially biologically important role during predictions. Also, this value is larger than the MEAN + *γ* STD threshold, defined for each PDB ID and each head.

The performance metrics for ERGO-II’s best model were obtained directly from their research paper repository. Their ROCAUC for McPAS and VDJDB were 0.939 and 0.866, respectively. However, they ceased weight updates with the use of the test set presumably for better use for the code repository users. This prevented us from replicating their top-performing model’s predictions accurately, thus hindering a fair comparison on the average precision.

### The models exhibit limited performance in the recent data test set

After confirmation of the model performance, we retrained the model with a larger dataset (herein referred to as the “entire data dataset”) consisting of McPAS and the whole VDJdb including the 10x Genomics dataset (10x Genomics, 2019)

Then we applied our entire-data-trained model to recently published data, the recent data test set, evaluating its efficacy in predicting TCR-peptide interactions in a real-life setting. Our objective with the entire data dataset approach was not necessarily to maximize generalizability but to uncover meaningful relationships and mimic the binding nature of the interactions. As shown in Table 4, most of the models did not achieve more than 0.9 ROCAUC scores, as in the benchmark, on the pure recent data dataset, as they did in the benchmarks. This result should be explained by the difficulties associated with the number of duplications; it is a difficult task if the interactions comprise the unseen CDRs or unseen peptides.

**Table 4.** Result of the recent data test dataset. APS stands for Average Precision Score.

| Model | Dataset | ROCAUC | APS | Number of interactions | Positive rate |
| --- | --- | --- | --- | --- | --- |
| Our model | Recent data test set | 0.5362 | 0.1855 | 33360 | 0.1667 |
|  | Recent data test set of new peptide subset | 0.5085 | 0.1707 | 28422 | 0.1662 |
|  | Recent data test set of known peptide subset | 0.6598 | 0.3318 | 4938 | 0.1692 |
|  | Recent data test set of new CDR3s subset | 0.5355 | 0.1844 | 33335 | 0.1660 |
| NetTCR2.0 | Recent data test set | 0.5274 | 0.1808 | 33360 | 0.1667 |
|  | Recent data test set of new peptide subset | 0.5113 | 0.1705 | 28422 | 0.1662 |
|  | Recent data test set of known peptide subset | 0.6327 | 0.3008 | 4938 | 0.1692 |
|  | Recent data test set of new CDR3s subset | 0.5267 | 0.1798 | 33335 | 0.1660 |
| PanPep * | Recent data test set | 0.5337 | 0.1897 | 30221 | 0.1745 |
|  | Recent data test set of new peptide subset | 0.5359 | 0.1908 | 25661 | 0.1739 |
|  | Recent data test set of known peptide subset | 0.5199 | 0.1852 | 4560 | 0.1779 |
|  | Recent data test set of new CDR3s subset | 0.5374 | 0.1923 | 29145 | 0.1752 |

The scores for the test set comprising only known CDR3s couldn't be computed as all the interactions are positive.
However, when setting a threshold at 0.5, our model achieves a recall score of 0.56, in comparison to NetTCR2.0's score of 0.44 and PanPep's 0.59.
\* The datasets employed in our model and NetTCR2.0 were identical. However, the dataset utilized in PanPep differed due to its exclusive use of CDR3 beta.
Consequently, by eliminating duplicates of the beta chain CDR3 from the test set, the total number of interactions was reduced from that of our model and NetTCR2.0.

After the training, the ROCAUC and the average precision score of the training dataset were 0.952 and 0.7952. Nonetheless, achieving generalizability against the recent data test set posed a significant challenge (Table 4), as evidenced by the ROCAUC and the average precision of the test set falling to 0.5362 and 0.1855 respectively. By restricting the interaction of the test set to the known peptide interactions, we did observe relative improvements in scores of average precision, rising to 0.3318. By restricting the interaction of the test set to the new peptide interactions that were not observed in the training dataset, we observed a deterioration of average precision to 0.1707. Not only does our model demonstrate poor performance, but the NetTCR2.0 or PanPep models also exhibit a similar level of performance deficiency in the recent data test set and its subsets. Regarding the PanPep model in which we used a zero-shot model setting for the unseen peptides and a majority model setting for the known peptides, while it claims to predict CDR3b-peptide pairs for unseen peptides, it achieved a slightly better average precision score of 0.1897, outperforming our model by a small margin. Nevertheless, for the interaction of the test data subset of the known peptide interactions, it only achieved less than the new peptide setting. These performances are still insufficient to serve as a viable alternative to wet lab experiments. Hence, it is clear that predicting CDR3-peptide interactions that contain peptides not represented in the training data remains a considerable challenge.

### Our model does not exhibit satisfactory performance for the Covid-19 dataset

We also applied our entire-data-trained model to a recently published Covid-19 dataset (Lu et al., 2021b), evaluating its efficacy in predicting TCR-peptide interactions in a real-life setting. As described in the Methods section, peptides from each SARS-CoV-2 protein were created with a 10-length residue window by moving one stride. No peptides of the Covid-19 dataset were found in the entire data dataset. The total number of interactions was 2,118,464, of which 2,118,434 were negative interactions and only 30 interactions were positive. In the 2,118,464 interactions, there were 1,676 unique CDR3 alpha-beta pairs and 1,264 unique peptides (1, 676 *·*1, 264 = 2, 118, 464, shown in Table 1). Out of the 30 positive interactions, we found 10 unique CDR3*α β* pairs and 18 unique peptides. Consequently, this means that the remaining 150 interactions, composed of these specific CDR3s and peptides, are classified as negative interactions (10 *·* 18 *−* 30 = 150).

By maximizing the precision of this prediction task, we adjusted a threshold to 0.9569, achieving a precision of 3.303*·*10^*−*5^, and a recall of 0.2333. In the confusion matrix, the True Positive count was 7, the False Negative count was 23, the False Positive count was 213,151, and the True Negative count was 1,905,283. The ROCAUC score was 0.4959 and the average precision score was 1.821*·* 10^*−*5^. Given the fact that positive interactions exist at a rate of 1.416 *·*10^*−*5^ (= 30*/*2118464), we can claim that the model can detect positive interactions 2.33 times (= 3.303*/*1.416) better than the random selection, but its specificity was not adequate enough to replace wet experiments.

### Residues in structural data are categorized based on their level of attention, into groups of largely attended and less attended ones

Although perfect generalizability wasn’t achieved, we sought to interpret the model within the 39 complex structures where the model surely performs well enough to analyze. Using the procedure described in the Methods section, we started with 47 TCR-related structures from the PDB Search and the SCEptRe server. Of these 47, our model designated 39 as having positive interactions, using a threshold of 0.5. A notable observation was that 30 out of these 39 structures share sequences with the entire data dataset. We paid special attention to these 39 cases in our analysis of attention layers, on the premise that the model’s accurate interpretability could be safely assumed for these instances. This is similar to a regression analysis examining the effect of some explanatory variables on target variables, and our goal was to identify the important features that the model learns, i.e., the features of the largely attended amino acid residues. We have provided details of these 39 structures in the Supplementary Information.

The attention values were considered “large” when they exceeded the threshold of MEAN + 5.5 STD on the peptide side and MEAN + 4.5 STD on the TCR side (5.5 and 4.5 are *γ*s in Equation 3). Approximately 20% of the residues were identified as large on each side, using *γ* as a result of the total sum of the four heads. The thresholds were determined through empirical evaluation, and the residue count generated by changing *γ* is provided in the Supplementary Information. The chosen thresholds were found to be effective in differentiating between large and small attention values. Note that the threshold for large attention values varies for each PDB entry or head due to differences in the distribution of attention values.

The analysis was performed separately for each of the four heads in the cross-attention layer (heads 0 to 3) on both the TCR and the peptide sides, with each head being analyzed separately. The cross-attention layer was defined on a CDR3*α β* sequence and a peptide sequence, resulting in an attention matrix with a shape determined by the length of the peptide and CDR3*α β* residues. It was possible for a particular residue to have a large attention value in head 0 but not in the other heads (as seen in Equation 3).

As an example, the attention values for the TCR-peptide complex of PDB entry 5TEZ are shown as eight heatmaps in Figure 4. The 5TEZ has the complex structure of MHC class I HLA-A2, influenza A virus, and TCRs (Yang et al., 2017). The corresponding 3D structure of the TCR-peptide complex is shown in Figure 5, in which the amino acid sequences of the peptide, CDR3*α*, and CDR3*β* are `GILGFVFTL`, CAASFIIQGAGKLVF, and CASSLLGGWSEAFF, respectively.

**Figure 5.**
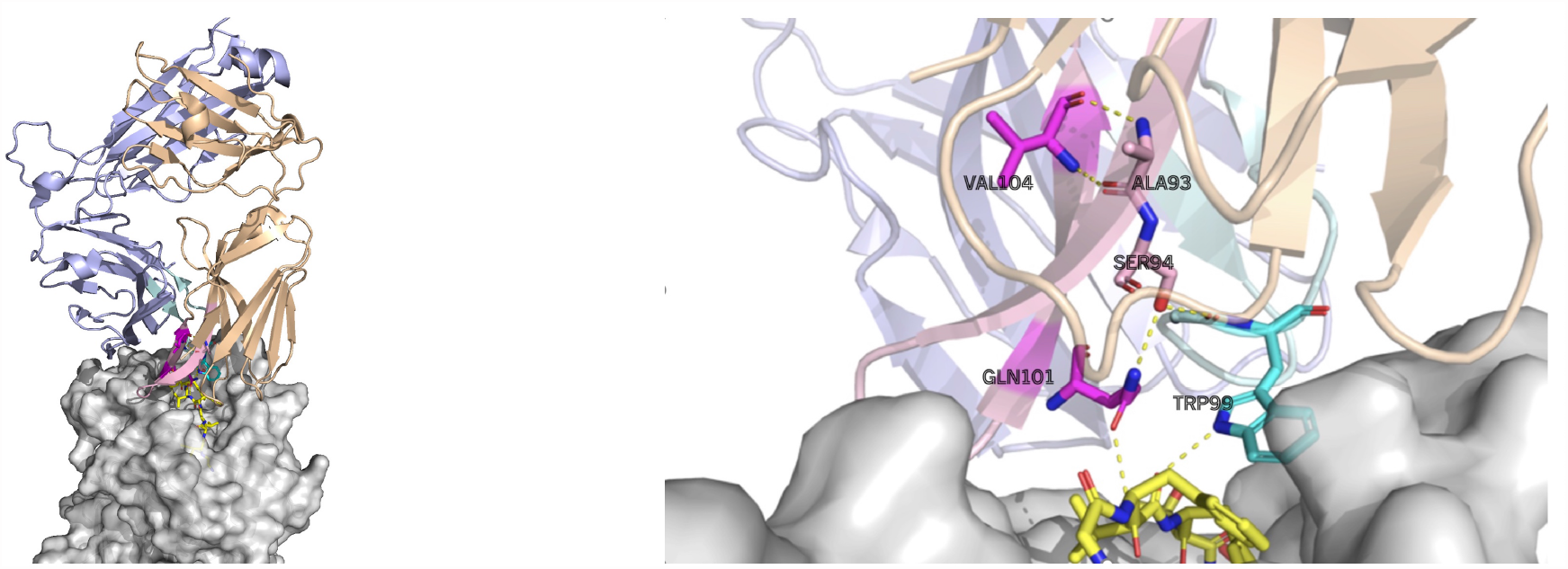
The largely attended residues in the TCR -the influenza virus epitope -HLA complex (PDB ID: 5TEZ), where the CDR3 sequences of TCR*α* and *β* are CAASFIIQGAQKLVF and CASSLLGGWSEAFF, respectively. The left figure shows the overall structure of the complex and the right figure shows the interactions of the largely attended residues: VAL104 (the 14th Val(V) of TCR*α*) and GLN101 (the 11th Gln(Q) of TCR*α*) of TCR *α* chain, and TRP99 (the 9th Trp(W)) of TCR*β* chain. The TCR *α* chain is wheat, the *β* chain is light-blue, the TCR*α* CDR3 part is light-pink, and the *β*’s CDR3 is pale-cyan. The residues with the large attention in the CDR3*α* are magenta, and that in the TCR*β* CDR3 is cyan. MHC is grey. The residues with the large attention and interacting residues are represented as sticks. The yellow dot lines represent the hydrogen bonds. VAL104 makes the two hydrogen bonds to TCR*α* ALA93 (the 3rd Alanine Ala(A) of TCR*α*) and may contribute to the stabilization of the end of the CDR3 loop conformation. GLN101 is hydrogen bonded with TCR*α* SER94, and SER94 is hydrogen bonded to the TCR*β*, maintaining the *α* and *β* structures. GLN101 of TCR*α* and the TRP99 of *β* have hydrogen bonds with the epitope. PyMol (Schrödinger and DeLano, 2020) is used for the visualization.

### Statistical analysis shows largely attended residues form H-Bonds with CDR3

Using the *γ* factor of 4.5 defined in Equation 3, we classified the TCR residues into two groups based on their attention values, “large” and “small,” for the cross-attention of the TCR side given a peptide. To gain insights into the characteristics of each group of residues, we analyzed their structural properties.

To assess differences between the two groups, we performed a paired t-test to remove variations arising from individual structural factors. In this study, 39 TCR-peptide complex structures were used as subjects, and structural properties associated with large or small attention groups were the tested values. The purpose of the paired t-test was to examine the null hypothesis that the mean difference between the pairs of measurements is zero. The proportion of a property, the test value (e.g. H-bonded to any peptide residue), is calculated by *P* = *A*_*h*_*/B*_*h*_, where *A*_*h*_ is the number of residues with one or more H-bonds of the specified type within the residues of large attention values, and *B*_*h*_ is the number of residues of large attention values, where *h* denotes the head.

The results of the statistical tests are shown in Table 5. Although each head was analyzed equally and separately, they showed different results. Head 0 happened to show the most significant differences, thus the results of head 0 and all concatenated heads are shown in Table 5. The results of the other heads are reported in the Supplementary Information.

**Table 5.** TCR-side attention analysis. Structural property comparisons between the large and small attention residue groups are shown. We showcase the results from head 0 and the results from all heads, as head 0 yielded the most significant findings.

| Property | Large Attention <sup>1</sup> | Small Attention <sup>1</sup> | p-value | Head |  |
| --- | --- | --- | --- | --- | --- |
| H-bonded to any peptide residue | 0.0256 ± 0.1581 | 0.0853 ± 0.0577 | 0.043 | 0 | *** |
| H-bonded to any CDR3 residue | 0.6068 ± 0.4262 | 0.4135 ± 0.1061 | 0.0111 | 0 | *** |
| H-bonded to any non-CDR3 TCR residue | 0.1966 ± 0.3734 | 0.4479 ± 0.0738 | 4.82e-04 | 0 | *** |
| H-bonded to any TCR residue | 0.7308 ± 0.3897 | 0.7193 ± 0.0903 | 0.864 | 0 |  |
| H-bonded to any CDR3 residue of own chain | 0.6068 ± 0.4262 | 0.3792 ± 0.0991 | 0.0027 | 0 | *** |
| H-bonded to any TCR residue of own chain | 0.7051 ± 0.4038 | 0.6408 ± 0.0888 | 0.353 | 0 |  |
| H-bonded to any CDR3 residue of opposite chain | 0.0000 ± 0.0000 | 0.0642 ± 0.0673 | 8.37e-07 | 0 | *** |
| In the edge <sup>2</sup> | 0.8376 ± 0.3235 | 0.5904 ± 0.0387 | 5.1e-05 | 0 | *** |
| Closest distance to peptide (Å) <sup>3</sup> | 9.1216 ± 3.1274 | 8.3528 ± 1.0097 | 0.119 | 0 |  |
| Number of H-bonds formed <sup>3</sup> | 1.7479 ± 1.1003 | 2.0919 ± 0.6806 | 0.0397 | 0 | *** |
| H-bonded to any peptide residue | 0.0862±0.1368 | 0.0805±0.0675 | 0.828 | all |  |
| H-bonded to any CDR3 residue | 0.4846±0.2216 | 0.4103±0.1040 | 0.0478 | all | *** |
| H-bonded to any non-CDR3 TCR residue | 0.2940±0.1923 | 0.4672±0.0846 | 3.88e-05 | all | *** |
| H-bonded to any TCR residue | 0.6845±0.1650 | 0.7294±0.0880 | 0.0987 | all |  |
| H-bonded to any CDR3 residue of own chain | 0.4643±0.2180 | 0.3752±0.0922 | 0.0107 | all | *** |
| H-bonded to any TCR residue of own chain | 0.6013±0.1999 | 0.6561±0.0880 | 0.117 | all |  |
| H-bonded to any CDR3 residue of opposite chain | 0.0306±0.0857 | 0.0672±0.0743 | 0.0369 | all | *** |
| In the edge <sup>2</sup> | 0.6434±0.2064 | 0.5928±0.0570 | 0.218 | all |  |
| Closest distance to peptide (Å) <sup>3</sup> | 8.4072±2.2892 | 8.4122±0.9592 | 0.988 | all |  |
| Number of H-bonds formed <sup>3</sup> | 2.0234±0.9370 | 2.0875±0.6685 | 0.589 | all |  |

1. Mean and standard deviation (for the 39 structures) of the proportion of residues that satisfy the property shown in the first column.
2. Four residues from the beginning and four from the end of the CDR. 3. In the last two properties, per-residue averages were used instead.

As a TCR sequence for the structural analysis includes both CDR3 and non-CDR3 portions, the H-bond properties were measured by dividing the residues into CDR3 and non-CDR3 portions.

The residues with large attention values had a more significant proportion of having an H-bond with the CDR3 portions inside their own chain. Nonetheless, the proportion of residues that are H-bonded to any TCR residue (i.e., H-bonds within the TCR chains) showed no difference between the large and small attention groups. This significant difference was observed both in head 0 and in all heads concatenated.

A natural consequence of those observations is that the largely attended residues are less likely to be H-bonded to the non-CDR3 portions, compared to the residues with small attention values. The most significant property difference in all concatenated heads occurred in the proportion of H-bonded to any non-CDR3 TCR residue. This means that, the largely attended residues are highly likely to avoid the H-bonded to the non-CDR3 TCR part, whereas they are likely to have H-bonds with the CDR3 portions.

In contrast, contrary to expectations, the proportion of largely attended TCR residues to form an H-bond with any peptide residues was not significant in all heads. Also, it is significantly smaller than that for small attended residues in head 0 with a p-value of 0.043. This highlighted a surprising and counter-intuitive finding in our analysis. We also examined the closest distance from a given TCR residue to any peptide residue, however, no significant difference was observed.

We also performed a similar analysis on the peptide side (Table 6) and observed that amino acid residues with large attention values of head 3 had significantly smaller distances to the closest TCR residues, a pattern not observed on the TCR side. Furthermore, the largely attended residues from all heads had shorter distances, though the difference was not statistically significant. This property was not observed in the TCR side attention and it poses an interesting structural aspect.

**Table 6.** Peptide side attention analysis. Structural property comparisons between the large and small attention residue groups are shown. We showcase the results from head 3 and the results from all heads, as head 3 yielded the most significant findings.

| Property | Large Attention <sup>1</sup> | Small Attention <sup>1</sup> | p-value | Head |  |
| --- | --- | --- | --- | --- | --- |
| H-bonded to any peptide residue | 0.1200±0.3250 | 0.0620±0.1108 | 0.31 | 3 |  |
| H-bonded to any CDR3 residue | 0.2800±0.4490 | 0.1714±0.0963 | 0.196 | 3 |  |
| H-bonded to any TCR residue | 0.4000±0.4899 | 0.2491±0.1120 | 0.0866 | 3 |  |
| In the edge <sup>2</sup> | 0.2600±0.4271 | 0.5771±0.1191 | 0.00591 | 3 | *** |
| Closest distance to TCR (Å) <sup>3</sup> | 3.7267±1.0947 | 5.1352±1.0994 | 5.15e-05 | 3 | *** |
| Number of H-bonds formed <sup>3</sup> | 1.7200±1.2496 | 2.0264±0.7809 | 0.503 | 3 |  |
| H-bonded to any peptide residue | 0.0495±0.1443 | 0.0659±0.1206 | 0.458 | all |  |
| H-bonded to any CDR3 residue | 0.2050±0.3024 | 0.1682±0.0982 | 0.48 | all |  |
| H-bonded to any TCR residue | 0.3401±0.3714 | 0.2372±0.1184 | 0.151 | all |  |
| In the edge <sup>2</sup> | 0.4459±0.4097 | 0.5874±0.1232 | 0.0795 | all |  |
| Closest distance to TCR (Å) <sup>3</sup> | 4.6398±1.7149 | 5.1926±1.2647 | 0.141 | all |  |
| Number of H-bonds formed <sup>3</sup> | 2.1126±1.4959 | 2.0031±0.9051 | 0.668 | all |  |

1. Mean and standard deviation (for the 39 structures) of the proportion of residues that satisfy the property shown in the first column.
2. Three residues from the beginning and three from the end of the peptide. 3. In the last two properties, per-residue averages were used instead.

### Impact of largely attended residues on model behaviors through input perturbation analysis

In this subsection, we delve into the effect of input perturbations on the outcomes of predictions and attention values through modification of the input sequence. This technique was utilized on the training data, PDBID 5TEZ. Furthermore, we extended this approach to a mutation study (Cole et al., 2013) which was not a part of our training data. The study involved mutating the protein sequence of the CDR3 beta loop of A6-TCR and assessing its binding strength against the TAX peptide, a peptide of the Human T-cell leukemia virus type I, on the MHC class I HLA-A2. The sequences and structures following mutation were recorded in the PDB under the identifiers, PDBID 1AO7 (before mutation) and PDBID 4FTV (after mutation).

For the 5TEZ PDB structure, three residues exhibited large attention values, 11th Gln(Q) of CDR3*α*, 14th Val(V) of CDR3*α*, and 9th Trp(W) of CDR3*β* (Table 7). We assessed how prediction and attention values were affected when these residues were substituted with alternative amino acids. The CDR3*α* sequence of 5TEZ is *CAASFIIQGAQKLVF*, while CDR3*β* is *CASSLLGGWSEAFF*. Notably, 11th Gln(Q), 14th Val(V), and 9th Trp(W) formed H-bonds but only the 14th Val of the *α* chain formed two H-bonds with the internal CDR3 chain of the TCR residue.

**Table 7.**
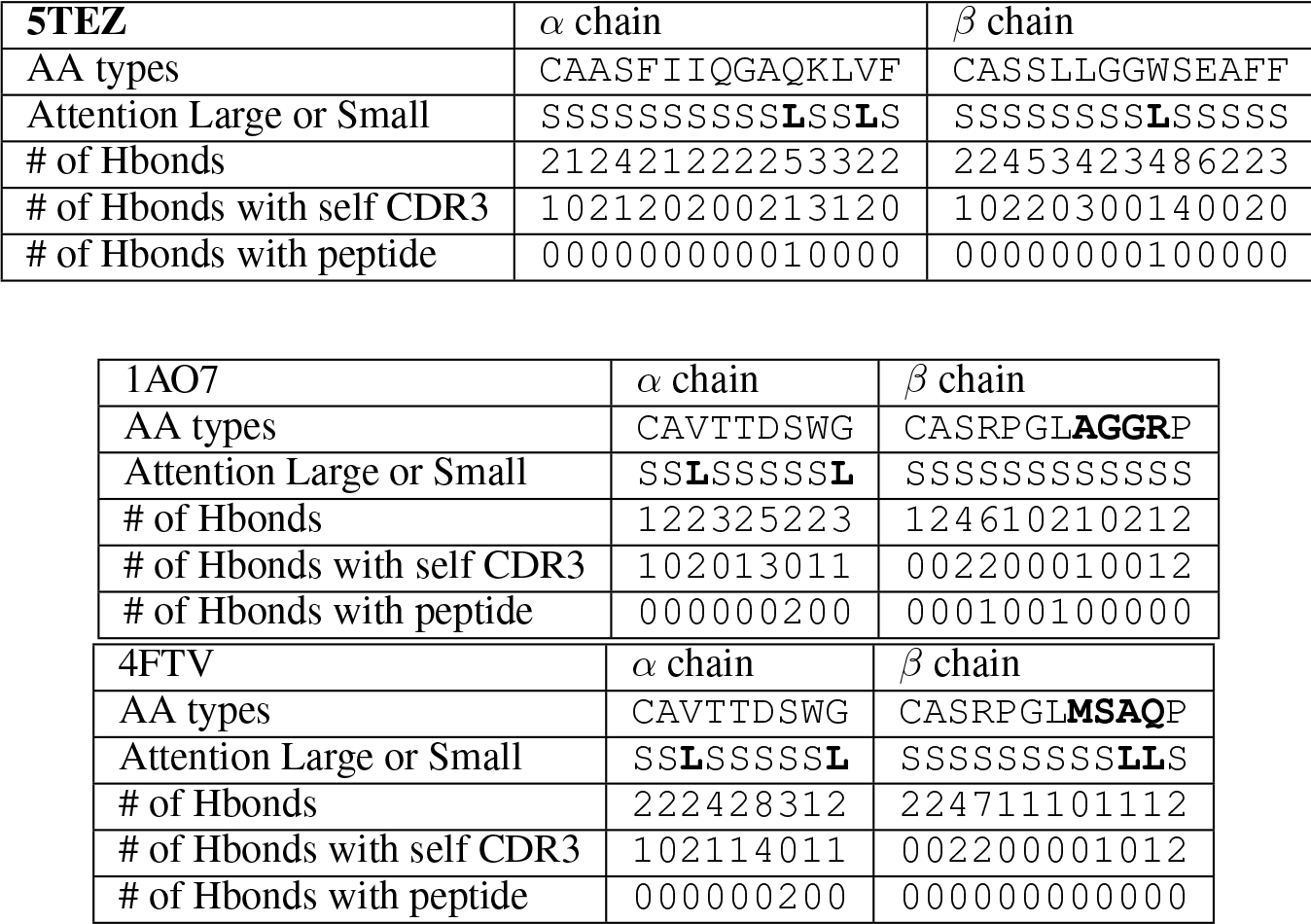
CDR3 chain analysis for 5TEZ, 1AO7 (Before mutation) and 4FTV (After mutation).

When the 14th Val(V) of the CDR3*α* was altered, the predictions experienced the most substantial impact, with the “unbound” prediction typically falling below 0.9 (Figure 6), probably because this attended residue has two HBonds with the CDR. Changes to the 11th Gln(Q) of the *α*, had a relatively minor effect on predictions, whereas alterations to the 9th Trp(W) of the *β* chain modify predictions while maintaining positive predictions with various amino acid substitutions. These results can be also confirmed by Figure 5 and these results support our hypothesis that the internal H-bonded structure of the CDR3 is crucial for peptide binding.

**Figure 6.**
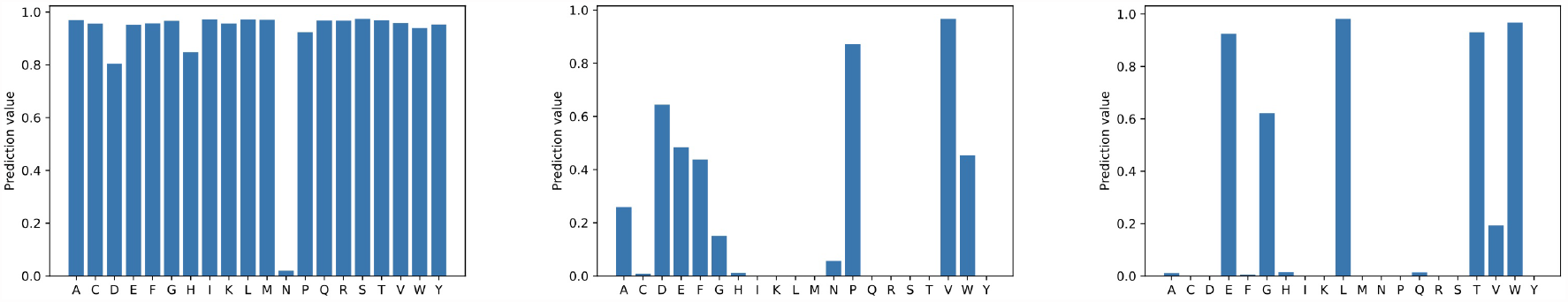
The first figure shows the prediction change of Glutamine(Q) of CDR3*α* from CAASFIIQGAQKLVF to CAASFIIQGA(-)KLVF. The center shows the prediction change of Valine(V) of CDR3*α* change: from CAASFIIQGAQKLVF to CAASFIIQGAQKL(-)F. The last shows the prediction change of Tryptophan(W) of CDR3*β*: from CASSLLGGWSEAFF to CASSLLGG(-)SEAFF. Notably, only the Valine(V) of CDR3*α* has two H-bonds with the internal CDR3 alpha chain, and hence, altering it has the most substantial impact.

In the 1AO7 (before mutation) and 4FTV (after mutation) structures, the previous study (Cole et al., 2013) identified that mutations in the four residues of the CDR3*β* chain of the TCR enhanced binding to the peptide nearly 1000-fold. We evaluated how predicted y-values change when these amino acid residues are substituted, focusing on the two structures with mutations. As shown in Table 7, the CDR3*α* sequence of 1AO7 and 4FTV is CAVTTDSWG, with CDR3*β* for 1AO7 being CASRPGLAGGRP and for 4FTV being CASRPGLMSAQP. The 4FTV mutation was from AGGR to MSQP, with the 8th to 11th residues enhancing affinity. Remarkably, our model successfully focused on the mutated residues in the 10th Ala(A) and 11th Gln(Q) of the *β* chain, although the model predicted both of them as positive. Furthermore, the study (Cole et al., 2013) posited that the mutation led to the loss of one hydrogen bond with the peptide, but the overall affinity was stronger after the mutation, suggesting an indirect contribution to the binding except for the TCR-peptide H-bonds. This finding also should reinforce our assertion that the internal H-bonded structure of the CDR3s is essential for peptide binding and reinforces the biological significance of attention values.

## DISCUSSION

### Interpretation of prediction in protein sequences

Prediction results of machine learning models are generally difficult to interpret, but when a model is used for binding predictions of biological sequences, the interpretability of the neural network model is essential. Highlighting residue positions is useful for understanding model predictions and is critical for utilizing later-stage applications. Our work, the Cross-TCR-Interpreter, tries to enable this interpretation with the attention layer.

Unexpectedly, we found that CDR residues with large attention values in our ML model were not necessarily interacting directly with peptide residues, as statistically shown in Table 5. Our result suggests that the ratio of hydrogen bond formation between CDR3s and a peptide can be relatively small, yet result in positive predictions in the model.

Instead, the TCR residues with large attention values appeared to stabilize a specific loop conformation required for the peptide binding, by forming H-bonds within the CDR3s. Accordingly, researchers have observed that residues that comprise the H-bond network within the TCR may be evolutionarily conserved (Garcia et al., 1996; Andrade et al., 2019), and the internal organization of the interface plays an important role in protein-protein interactions (Rauf et al., 2009; Reichmann et al., 2005). Our findings suggest that certain residues may be oriented in a specific direction with internal H-bonds and the attention layer may emphasize their importance in TCRs in terms of binding stability.

Our findings also showed that the average distance between the TCR and peptide in 3D decreased when the attention value on the peptide side of attention was large. This may be because the peptide, being a short sequence, has a limited contribution to TCR binding that is related solely to distance. However, the larger attention values on the TCR side did not necessarily correspond to smaller distances in the 3D structure, potentially because TCRs are longer and more complicated in their binding role.

Not all the sequence paired data were available with 3D structure and we knew that the number was small, but we experimented with as many available structures as possible. We used the 3D structure as the confirmation of attention layer interpretation. In future investigations, it may be possible to use a different machine learning model such as a meaningful perturbation method on exhaustively collected sequences.

### Model limitations due to the dataset and difficulties associated with the recent data and Covid-19 dataset

Our analysis and results might, admittedly, raise several questions regarding the interpretability of attention values observed between TCR and pMHC. The diverse sampling of TCR in comparison to the peptide samples in our dataset might influence the apparent association of significant attention with structural properties, such as hydrogen bond formation, primarily within the TCR. This imbalance could help explain why our model predominantly associates the presence of certain TCR residues with reactivity against a specific peptide, rather than assigning weight to peptide-side attention.

The approach to generating negative data in benchmark datasets may skew our test dataset to resemble our training data more than would be typical in real-world prospective evaluation scenarios. This is indicated by the inferior performance of the recent data test set scores and the Covid-19 dataset results. The difficulty of prediction in the recent data test set and the Covid-19 data was not caused by differences in TCR but instead by differences in peptides between the Covid-19 data and baseline data. We plotted the sequence-sequence pairwise distance matrix with UMAP dimension reduction, as shown in Figure 7. There was no difference in the distribution of TCR between the entire data dataset data and the Covid-19 data, whereas there was a substantial difference in the distribution of peptides. This discussion is also supported by previous studies on TCR predictions (Weber et al., 2021; Moris et al., 2021; Essaghir et al., 2022), in which the authors stated that generalization and extrapolation to unseen epitopes remain challenging.

**Figure 7.**
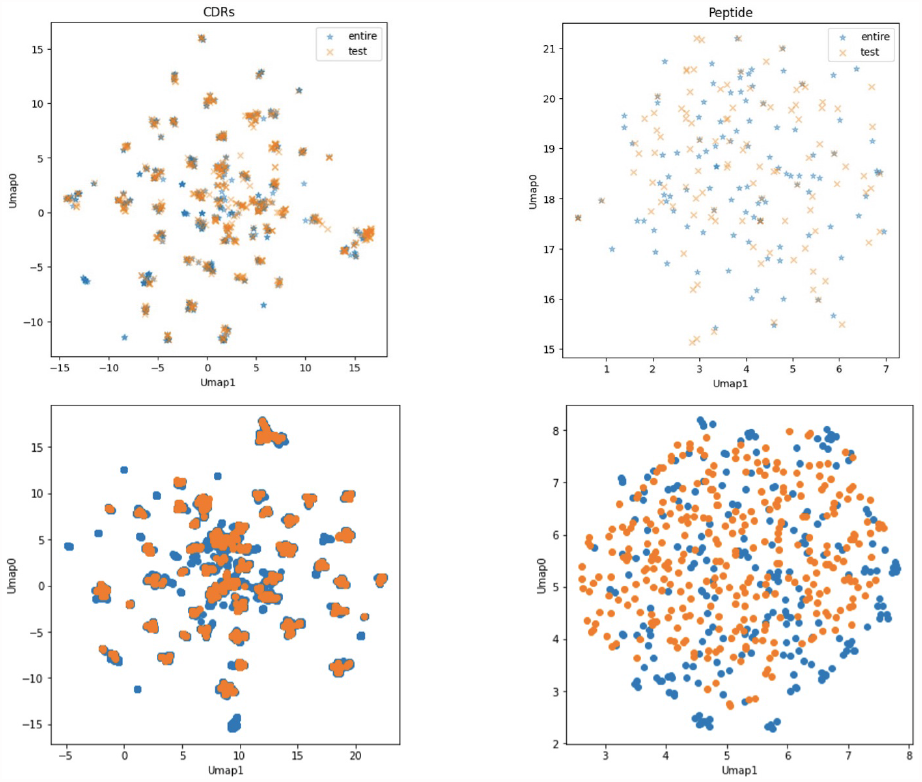
UMAP visualizations of sequence distance maps of CDR3s and peptides. Each point shows a sequence datum. The two colors on the peptide side (right) show minimal overlap, indicating different peptides of the entire data dataset. Upper half: the recent data test set. Lower half: the Covid-19 dataset. Left two figures: sequences of CDR3*α β* visualization. Right two figures: peptide sequences visualization. Orange points: the recent data test set (Upper) and the Covid-19 dataset (Lower). Blue points: the entire data dataset.

In addition, when the Covid-19 dataset was modified to a positive ratio of 20%, the model’s ROCAUC value was 0.5692 and the average precision score was 0.3312. When we set the threshold to 0.9977, the precision was 0.6667 and the recall was 0.06667. The positive ratio affected the performance of the model evaluation.

### Difficulties associated with unseen data

Additionally, we sought to evaluate the performance of our model specifically on interactions involving unseen peptides or different TCRs within the benchmark test set of McPAS and VDJDB data. This was done either by removing the interactions of peptides of the training dataset or by removing similar TCRs from the test dataset.

Although the majority of the peptides were already present in the training data, we identified a subset of 46 interactions (14 positives) for McPAS and 16 interactions (8 positives) for VDJDB that involved unseen peptides (the numbers, 46 and 16, are also indicated in Figure 2). The ROCAUC scores for these unseen peptide interactions were 0.721 for McPAS and 0.719 for VDJDB, which were a lot lower than the scores reported in Table 2 and Table 3 for interactions involving seen peptides. This performance gain when evaluating the model on already seen peptides was also observed in the recent data test set experiment.

In Figure 8, our performance metrics indicated a decline when we refined the test dataset by eliminating any test interactions involving TCRs that exhibit a distance greater than a certain threshold value from the TCRs present in the training set. This trend underscores the sensitivity of the model to the diversity and distribution of TCRs in the test data.

**Figure 8.**
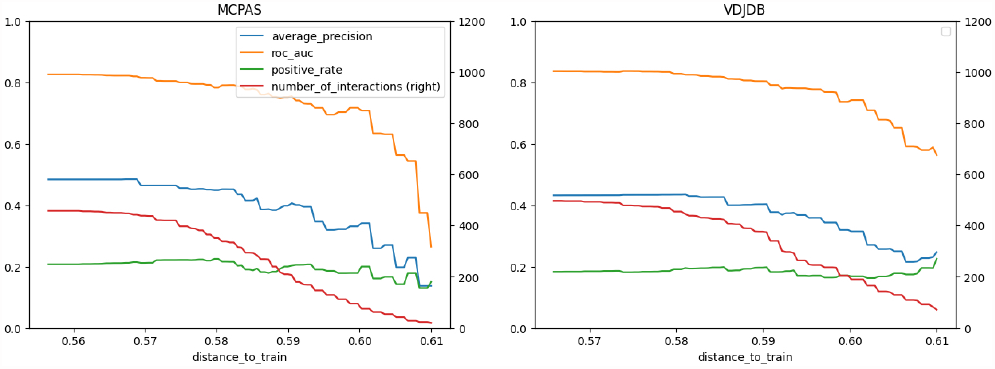
Performance delegation of the peptide `GILGFVFTL` by removing similar TCRs interactions from the test set. The x-axis shows the distance threshold. For instance, if the x-value is 0.58, no test data less than that distance is used for evaluation.

## CONCLUSION

Our study presents a computational approach for predicting T cell receptor (TCR) binding to specific ligand peptides. Our study predicted TCR-peptide binding with the cross-attention mechanism as well as analyzed the available protein structures comprehensively to gain new insights into TCR-peptide functional relationships.

By incorporating an attention layer based on language models, our machine learning model achieved competitive performance on a benchmark dataset of TCR-peptide binding, although it confronted enduring challenges with the Covid-19 dataset and the recent data test set.

Our analysis of the model allowed us to associate neural network weights with protein 3D structure datasets, identify statistically significant properties of largely attended residues, and detail the binding principle through the visualization and analysis of the cross-attention layer, source-target-attention layer.

The statistical analysis of the attention layer on the structural data revealed that the largely attended residues were more likely to contact their own CDR3 than normal residues, thereby providing new insights into the TCR-peptide binding mechanisms. Proteins create hydrogen bonds to form special structures and may play special roles when a peptide is conditioned to react with them.

## AUTHOR CONTRIBUTIONS

Kyohei Koyama developed the model, analyzed the results, and wrote the manuscript as part of his doctoral work. Kosuke Hashimoto, Chioko Nagao, and Kenji Mizuguchi advised and Kenji Mizuguchi managed the entire project. All authors have contributed to the manuscript and approved the submitted version.

## ACKNOWLEDGMENTS

We wish to thank the lab members for their supportive discussion.

## DATA AVAILABILITY STATEMENT

Code and data are available at https://github.com/kyoheikoyama/TCRPrediction

## REFERENCES

10x Genomics (2019). A New Way of Exploring Immunity–Linking Highly Multiplexed Antigen Recognition to Immune Repertoire and Phenotype. Tech. rep

Akiba, T., Sano, S., Yanase, T., Ohta, T., and Koyama, M. (2019). Optuna: A next-generation hyperparameter optimization framework. In Proceedings of the 25th ACM SIGKDD international conference on knowledge discovery & data mining. 2623–2631

Andrade, M., Pontes, C., and Treptow, W. (2019). Coevolutive, evolutive and stochastic information in protein-protein interactions. Computational and Structural Biotechnology Journal 17, 1429–1435

Berman, H., Henrick, K., and Nakamura, H. (2003). Announcing the worldwide protein data bank. Nature Structural & Molecular Biology 10, 980–980

Chapman, B. and Chang, J. (2000). Biopython: Python tools for computational biology. ACM Sigbio Newsletter 20, 15–19

Chen, J., Yang, L., Raman, K., Bendersky, M., Yeh, J.-J., Zhou, Y., et al. (2020). DiPair: Fast and accurate distillation for trillion-scale text matching and pair modeling. arXiv preprint arXiv:2010.03099

Cole, D. K., Sami, M., Scott, D. R., Rizkallah, P. J., Borbulevych, O. Y., Todorov, P. T., et al. (2013). Increased peptide contacts govern high affinity binding of a modified TCR whilst maintaining a native pMHC docking mode. Frontiers in immunology 4, 168

Dash, P., Fiore-Gartland, A. J., Hertz, T., Wang, G. C., Sharma, S., Souquette, A., et al. (2017). Quantifiable predictive features define epitope-specific T cell receptor repertoires. Nature 547, 89–93

Devlin, J., Chang, M.-W., Lee, K., and Toutanova, K. (2018). Bert: Pre-training of deep bidirectional transformers for language understanding. arXiv preprint arXiv:1810.04805

Dunbar, J. and Deane, C. M. (2016). ANARCI: antigen receptor numbering and receptor classification. Bioinformatics 32, 298–300

Essaghir, A., Sathiyamoorthy, N. K., Smyth, P., Postelnicu, A., Ghiviriga, S., Ghita, A., et al. (2022). T-cell receptor specific protein language model for prediction and interpretation of epitope binding (ProtLM. TCR). bioRxiv, 2022–11

Gao, Y., Gao, Y., Fan, Y., Zhu, C., Wei, Z., Zhou, C., et al. (2023). Pan-Peptide Meta Learning for T-cell receptor–antigen binding recognition. Nature Machine Intelligence, 1–14

Garcia, K. C., Degano, M., Stanfield, R. L., Brunmark, A., Jackson, M. R., Peterson, P. A., et al. (1996). An α β T cell receptor structure at 2.5 Å and its orientation in the TCR-MHC complex. Science 274, 209–219

Gheini, M., Ren, X., and May, J. (2021). Cross-Attention is All You Need: Adapting Pretrained Transformers for Machine Translation.In Proceedings of the 2021 Conference on Empirical Methods in Natural Language Processing. 1754–1765. doi:10.18653/v1/2021.emnlp-main.132

Gowthaman, R. and Pierce, B. G. (2019). TCR3d: The T cell receptor structural repertoire database. Bioinformatics 35, 5323–5325

Hao, Y., Dong, L., Wei, F., and Xu, K. (2021). Self-attention attribution: Interpreting information interactions inside transformer. Proceedings of the AAAI Conference on Artificial Intelligence 35, 12963–12971

Honda, S., Koyama, K., and Kotaro, K. (2020). Cross Attentive Antibody-Antigen Interaction Prediction with Multi-task Learning. ICML 2020 Workshop on Computational Biology (WCB)

Koyama, K., Kamiya, K., and Shimada, K. (2020). Cross Attention DTI: Drug-Target Interaction Prediction with Cross Attention module in the Blind Evaluation Setup. BIOKDD2020

Lee, K.-H., Chen, X., Hua, G., Hu, H., and He, X. (2018). Stacked cross attention for image-text matching. In Proceedings of the European conference on computer vision (ECCV). 201–216

Lu, T., Zhang, Z., Zhu, J., Wang, Y., Jiang, P., Xiao, X., et al. (2021a). Deep learning-based prediction of the T cell receptor–antigen binding specificity. Nature machine intelligence 3, 864–875

Lu, X., Hosono, Y., Nagae, M., Ishizuka, S., Ishikawa, E., Motooka, D., et al. (2021b). Identification of conserved SARS-CoV-2 spike epitopes that expand public cTfh clonotypes in mild COVID-19 patientsSARS-CoV-2 spike epitopes for public cTfh cells. Journal of Experimental Medicine 218. doi:10.1084/jem.20211327.E20211327

Mahajan, S., Yan, Z., Jespersen, M. C., Jensen, K. K., Marcatili, P., Nielsen, M., et al. (2019). Benchmark datasets of immune receptor-epitope structural complexes. BMC bioinformatics 20, 1–7

Montemurro, A., Schuster, V., Povlsen, H. R., Bentzen, A. K., Jurtz, V., Chronister, W. D., et al. (2021). NetTCR-2.0 enables accurate prediction of TCR-peptide binding by using paired TCRα and β sequence data. Communications Biology 4, 1–13

Moris, P., De Pauw, J., Postovskaya, A., Gielis, S., De Neuter, N., Bittremieux, W., et al. (2021). Current challenges for unseen-epitope TCR interaction prediction and a new perspective derived from image classification. Briefings in Bioinformatics 22, bbaa318

Parthasarathy, S. and Sundaram, S. (2021). Detecting expressions with multimodal transformers. In 2021 IEEE Spoken Language Technology Workshop (SLT) (IEEE), 636–643

Rauf, S. M. A., Ismael, M., Sahu, K. K., Suzuki, A., Sahnoun, R., Koyama, M., et al. (2009). A graph theoretical approach to the effect of mutation on the flexibility of the DNA binding domain of p53 protein. Chemical Papers 63, 654–661

Reichmann, D., Rahat, O., Albeck, S., Meged, R., Dym, O., and Schreiber, G. (2005). The modular architecture of protein-protein binding interfaces. Proceedings of the National Academy of Sciences 102, 57–62

Rogers, A., Kovaleva, O., and Rumshisky, A. (2020). A primer in bertology: What we know about how bert works. Transactions of the Association for Computational Linguistics 8, 842–866

[Dataset] Schrö dinger, L. and DeLano, W. (2020). PyMOL

Shugay, M., Bagaev, D. V., Zvyagin, I. V., Vroomans, R. M., Crawford, J. C., Dolton, G., et al. (2018). VDJdb: a curated database of T-cell receptor sequences with known antigen specificity. Nucleic acids research 46, D419–D427

Sidhom, J.-W., Larman, H. B., Pardoll, D. M., and Baras, A. S. (2021). DeepTCR is a deep learning framework for revealing sequence concepts within T-cell repertoires. Nature communications 12, 1–12

Sievers, F., Wilm, A., Dineen, D., Gibson, T. J., Karplus, K., Li, W., et al. (2011). Fast, scalable generation of high-quality protein multiple sequence alignments using Clustal Omega. Molecular systems biology 7, 539

Springer, I., Besser, H., Tickotsky-Moskovitz, N., Dvorkin, S., and Louzoun, Y. (2020). Prediction of specific TCR-peptide binding from large dictionaries of TCR-peptide pairs. Frontiers in immunology, 1803

Springer, I., Tickotsky, N., and Louzoun, Y. (2021). Contribution of T Cell Receptor Alpha and Beta CDR3, MHC Typing, V and J Genes to Peptide Binding Prediction. Frontiers in Immunology 12

Tickotsky, N., Sagiv, T., Prilusky, J., Shifrut, E., and Friedman, N. (2017). McPAS-TCR: a manually curated catalogue of pathology-associated T cell receptor sequences. Bioinformatics 33, 2924–2929

Vaswani, A., Shazeer, N., Parmar, N., Uszkoreit, J., Jones, L., Kaiser, Ł., et al. (2017). Attention is all you need. Advances in neural information processing systems 30

Voita, E., Talbot, D., Moiseev, F., Sennrich, R., and Titov, I. (2019). Analyzing Multi-Head Self-Attention: Specialized Heads Do the Heavy Lifting, the Rest Can Be Pruned. In Proceedings of the 57th Annual Meeting of the Association for Computational Linguistics (Florence, Italy: Association for Computational Linguistics), 5797–5808. doi:10.18653/v1/P19-1580

Wallace, A. C., Laskowski, R. A., and Thornton, J. M. (1995). LIGPLOT: a program to generate schematic diagrams of protein-ligand interactions. Protein engineering, design and selection 8, 127–134

Weber, A., Born, J., and Rodriguez Martínez, M. (2021). TITAN: T-cell receptor specificity prediction with bimodal attention networks. Bioinformatics 37, i237–i244

Wu, K., Yost, K. E., Daniel, B., Belk, J. A., Xia, Y., Egawa, T., et al. (2021). TCR-BERT: learning the grammar of T-cell receptors for flexible antigen-xbinding analyses. bioRxiv, 2021.11.18.469186

Xu, Y., Qian, X., Tong, Y., Li, F., Wang, K., Zhang, X., et al. (2022). AttnTAP: An attention-fused BiLSTM model used to predict TCRpeptide binding accuracy. Frontiers in Genetics, 1871

Xu, Z., Luo, M., Lin, W., Xue, G., Wang, P., Jin, X., et al. (2021). DLpTCR: an ensemble deep learning framework for predicting immunogenic peptide recognized by T cell receptor. Briefings in Bioinformatics 22, bbab335

Yang, X., Chen, G., Weng, N.-p., and Mariuzza, R. A. (2017). Structural basis for clonal diversity of the human T-cell response to a dominant influenza virus epitope. Journal of Biological Chemistry 292, 18618–18627. doi:10.2210/pdb5tez/pdb

