## Supplemental file for "Attention network for predicting T cell receptor-peptide binding can associate attention with interpretable protein structural properties"

### Supplementary Material

#### HYPERPARAMETERS OF THE MODEL

The parameter of the model was optimized by using a hyperparameter optimization package, Optuna. The best parameters are $batch\_size$ = 2949, $d\_ff$ of the final MLP layer = 84, $dim$ in the model = 256, $dropout\_rate$ = 7.651e-05, and $learning\_rate$ = 9.387e-05.

The number of transformer layers for self-attention is two on each side of the TCR and the peptide. The $n\_head$ of the transformer encoder is four. The number of cross-attention layers is one. The maximum length of TCR$\alpha$ and TCR$\beta$ in the dataset was 62, and that of the peptide was 26. Although an intensive and exhaustive search may have different results, initial searches on pre-training and transfer learning did not contribute to improving the final score.

#### COMPARISON OF OCCURRENCE OF AMINO ACID TYPES OF RESIDUES IN THE LARGE AND SMALL ATTENTION GROUPS

In general, TCRs have, in order of abundance, serine(S), alanine (A), and phenylalanine (F). On the other hand, TCRs with large attention values have valine (V), glycine (G), and glutamic acid (E). As the right panel of Figure S1 shows, methionine (M), valine (V), and lysine (K) tend to have relatively large attention values. The side chains of M and V are known as polarity-free and belong to the aliphatic group.

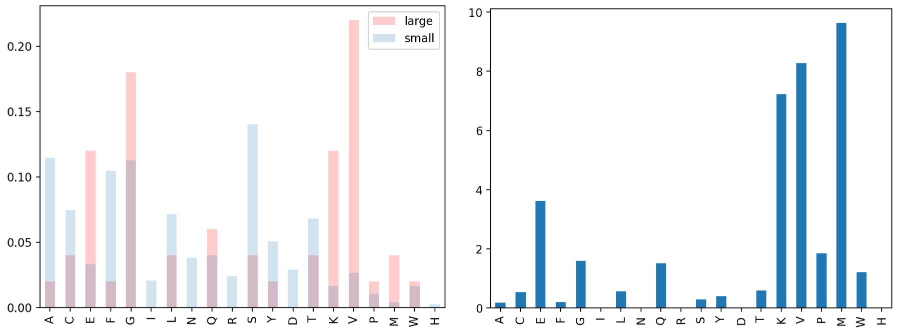


Figure S1: Left: Residue type distribution of large attention TCR group and small attention TCR group. Right: the value of the large group divided by the value of the small group, indicating the occurrence tendency of large attention value.

### INFULUENCE OF THE CHANGE OF THE $\gamma$ FACTOR

The attention values were considered ”large” when they exceeded the threshold of MEAN + 5.5 STD on the peptide side and MEAN + 4.5 STD on the TCR side (4.5 and 5.5 are $\gamma$ factors). Approximately 20% of the residues were identified as large on each side, using $\gamma$ as a result of the total sum of the four heads Table S1.

**Table S1.** Count of large or small attention residue groups when changing the factor $\gamma$. Four heads are merged.

| | **TCRs Large Attention** | **TCRs Small Attention** | **Peptides Large Attention** | **Peptides Small Attention** |
|---|---|---|---|---|
| **count ($\gamma$=2)** | 569 | 468 | 372 | 65 |
| **count ($\gamma$=2.5)** | 441 | 596 | 325 | 112 |
| **count ($\gamma$=3))** | 364 | 673 | 264 | 173 |
| **count ($\gamma$=3.5** | 299 | 738 | 219 | 218 |
| **count ($\gamma$=4))** | 244 | 793 | 172 | 265 |
| **count ($\gamma$=4.5** | 196 | 841 | 137 | 300 |
| **count ($\gamma$=5))** | 153 | 884 | 111 | 326 |
| **count ($\gamma$=5.5** | 118 | 919 | 81 | 356 |
| **count ($\gamma$=6))** | 97 | 940 | 48 | 389 |

### ADDITIONAL DATASET STATISTICS

Unique count statistics are shown in Table S2. The interaction column means the unique count of pairs of $\{$CDR3$\alpha$, CDR3$\beta$, Peptide$\}$ and CDR3$\alpha\beta$ denotes the unique count of pairs of $\{$CDR3$\alpha$, CDR3$\beta\}$. The duplication count means the number of unique data that is shared between training and test sets.

**Table S2.** Unique count statistics.

| Dataset name | CDR3$\alpha$ | CDR3$\beta$ | Interaction |
|---|---|---|---|
| McPAS, training | 2423 | 2560 | 23363 |
| McPAS, test | 718 | 714 | 4729 |
| McPAS, duplication count | 218 | 198 | 0 |
| VDJdb-without10x, training | 2151 | 2171 | 19526 |
| VDJdb-without10x, test | 570 | 572 | 4010 |
| VDJdb, duplication count | 198 | 196 | 0 |
| The entire data dataset | 17954 | 19162 | 119046 |
| The recent data test set | 4782 | 5174 | 33360 |
| The duplication count of the two above | 444 | 148 | 0 |

### ATTENTION CAN CAPTURE THE SIGN OF POSITIVE BINDING

In a minor experiment, we tested the attention mechanism of our model and its effectiveness using mock data. We added two Alanine (A) residues to the end of TCR alpha for every positive pair, indicating a clear and obvious sign of positive binding. After training the model on this mock data from McPAS, it achieved the ROCAUC score of 1.0. Upon visualizing the attention layer of the trained model, we found that the sign of positive interactions was indeed given large attention values, as shown in Figure S2. Hence, we used attention values as a tool to interpret the binding mechanism.

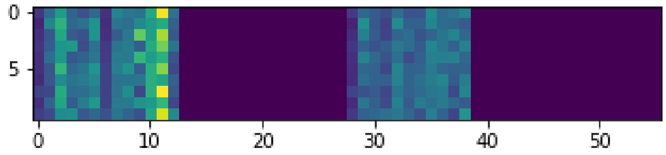


Figure S2: Attention values visualization on one of the mock dataset pairs. The position of the 11th residue was the first Alanine added to the TCR alpha chain, and it had large attention values. X-axis: Residues of TCRs. Y-axis: Residues of a peptide. Dark blue indicates zero attention values (padding) and yellow indicates large values.

### PDB IDS FOR TCR-PEPTIDE CRYSTAL

There are 39 PDB IDs for the analysis out of the 80 TCR-peptide crystal structures of PDB. 9 PDB IDs were not included in the entire data, whereas 30 PDB IDs were included.

- 55 PDBIDs before processing:

- 2VLK, 2ICW, 5WKF, 2VLJ, 3PQY, 4MJI, 4P2Q, 2YPL, 5BRZ, 6RPB, 1J8H, 4P2R, 5MEN, 3MV8, 4OZF, 3VXR, 3VXS, 4OZG, 3MV9, 5TEZ, 2J8U, 6Q3S, 4JRX, 5WLG, 3VXU, 1U3H, 4JRY, 4Z7V, 2UWE, 1LP9, 3W0W, 6AVF, 2BNQ, 4JFE, 4JFD, 3QIU, 6AVG, 2Z31, 2BNR, 5ISZ, 5KS9, 3MV7, 3MBE, 4OZH, 2NX5, 5NHT, 4QOK, 5D2L, 1D9K, 1FYT, 4P2O, 5WKH, 6EQB, 2VLR, 6EQA

- 47 PDBIDs having no identical sequences:

- 2VLK, 2ICW, 5WKF, 3PQY, 4MJI, 4P2Q, 2YPL, 5BRZ, 6RPB, 1J8H, 4P2R, 5MEN, 3MV8, 4OZF, 3VXR, 3VXS, 4OZG, 5TEZ, 2J8U, 6Q3S, 4JRX, 5WLG, 3VXU, 1U3H, 4JRY, 4Z7V, 6AVF, 4JFE, 4JFD, 3QIU, 6AVG, 2Z31, 2BNR, 5ISZ, 5KS9, 3MBE, 4OZH, 2NX5, 5NHT, 4QOK, 5D2L, 1D9K, 4P2O, 5WKH, 6EQB, 2VLR, 6EQA

- 39 PDB IDs having positive predictions:

- 2VLK, 5WKF, 3PQY, 4MJI, 4P2Q, 2YPL, 1J8H, 4P2R, 5MEN, 3MV8, 4OZF, 3VXR, 3VXS, 4OZG, 5TEZ, 2J8U, 6Q3S, 4JRX, 3VXU, 1U3H, 4JRY, 4Z7V, 4JFE, 4JFD, 3QIU, 2Z31, 2BNR, 3MBE, 4OZH, 2NX5, 5NHT, 4QOK, 5D2L, 1D9K, 4P2O, 5WKH, 6EQB, 2VLR, 6EQA

- 9 PDB IDs that are not in the entire sequence dataset:

- 5MEN, 2J8U, 3VXU, 4JFE, 4JFD, 5NHT, 4QOK, 6EQB, 6EQA

### S PROTEIN SEQUENCE OF COVID-19

- mfvflvllpl vssqcvnltt rtqlppaytn sftrgvyypd kvfrssvlhs tqdlflpffs nvtwfhaihv sgtngtkrfd npvlpfndgv yfasteksni irgwifgttl dsktqslliv nnatnvvikv cefqfcndpf lgvyyhknnk swmesefrvy ssannctfey vsqpflmdle gkqgnfknlr efvfknidgy fkiyskhtpi nlvrdlpqgf saleplvdlp iginitrfqt llalhrsylt pgdsssgwta gaaayyvgyl qprtfllkyn engtitdavd caldplsetk ctlksftvek giyqtsnfrv qptesivrfp nitnlcpfge vfnatrfasv yawnrkrisn cvadysvlyn sasfstfkcy gvsptklndl cftnvyadsf virgdevrqi apgqtgkiad ynyklpddft gcviawnsnn ldskvggnyn ylyrlfrksn lkpferdist eiyqagstpc ngvegfncyf plqsygfqpt ngvgyqpyrv vvlsfellha patvcgpkks tnlvknkcvn fnfngltgtg vltesnkkfl pfqqfgrdia dttdavrdpq tleilditpc sfggvsvitp gtntsnqvav lyqdvnctev pvaihadqlt ptwrvystgs nvfqtragcl igaehvnnsy ecdipigagi casyqtqtns prrarsvasq siiaytmslg aensvaysnn siaiptnfti svtteilpvs mtktsvdctm yicgdstecs nlllqygsfc tqlnraltgi aveqdkntqe vfaqvkqiyk tppikdfggf nfsqilpdps kpskrsfied llfnkvtlad agfikqygdc lgdiaardli caqkfngltv lpplltdemi aqytsallag titsgwtfga gaalqipfam qmayrfngig vtqnvlyenq klianqfnsa igkiqdslss tasalgklqd vvnqnaqaln tlvkqlssnf gaissvlndi lsrldkveae vqidrlitgr lqslqtyvtq qliraaeira sanlaatkms ecvlgqskrv dfcgkgyhlm sfpqsaphgv vflhvtyvpa qeknfttapa ichdgkahfp regvfvsngt hwfvtqrnfy epqiittdnt fvsgncdvvi givnntvydp lqpeldsfke eldkyfknht spdvdlgdis ginasvvniq keidrlneva knlneslidl qelgkyeqyi kwpwyiwlgf iagliaivmv timlccmtsc csclkgccsc gscckfdedd sepvlkgvkl

### STATISTICAL TEST RESULTS OF STRUCTURES USING DIFFERENT ATTENTION HEADS

The results of the paired t-tests are shown. Although each head was analyzed equally and separately, they showed different results (Table S3, Table S4).

**Table S3.** Peptide side attention analysis on all heads. In a cell, the left number is the mean value over the PDBIDs and the right number is the STD value.

| Property | Large Attention [1] | Small Attention [1] | p-value | Head | |
|---|---|---|---|---|---|
| Closest distance to peptide (Å) | 4.1233±1.3970 | 5.0796±1.1194 | 0.0703 | 0 | |
| Number of H-bonds formed | 1.6667±1.5986 | 2.0125±0.7579 | 0.84 | 0 | |
| H-bonded to any CDR residue | 0.0833±0.2764 | 0.1801±0.1086 | 0.409 | 0 | |
| H-bonded to any peptide residue | 0.0833±0.2764 | 0.0656±0.1218 | 1 | 0 | |
| H-bonded to any TCR residue | 0.3333±0.4714 | 0.2559±0.1198 | 0.552 | 0 | |
| H-bonded to any non-CDR TCR residue | 0.2500±0.4330 | 0.1167±0.1116 | 0.318 | 0 | |
| In the edge [2] | 0.2500±0.4330 | 0.5637±0.0951 | 0.0504 | 0 | |
| Closest distance to peptide (Å) | 5.1794±2.4525 | 5.0293±1.0674 | 0.895 | 1 | |
| Number of H-bonds formed | 2.4348±1.7087 | 1.9672±0.7934 | 0.154 | 1 | |
| H-bonded to any CDR residue | 0.3261±0.4570 | 0.1710±0.0996 | 0.215 | 1 | |
| H-bonded to any peptide residue | 0.0000±0.0000 | 0.0673±0.1226 | 0.0711 | 1 | |
| H-bonded to any TCR residue | 0.4130±0.4812 | 0.2499±0.1138 | 0.168 | 1 | |
| H-bonded to any non-CDR TCR residue | 0.1739±0.3790 | 0.1162±0.1197 | 0.457 | 1 | |
| In the edge [2] | 0.4783±0.4995 | 0.5610±0.1018 | 0.566 | 1 | |
| Closest distance to peptide (Å) | 4.5241±1.8359 | 5.0844±1.0686 | 0.0771 | 2 | |
| Number of H-bonds formed | 1.6304±1.0448 | 2.0183±0.7681 | 0.478 | 2 | |
| H-bonded to any CDR residue | 0.1522±0.3437 | 0.1791±0.1188 | 0.653 | 2 | |
| H-bonded to any peptide residue | 0.0435±0.2039 | 0.0659±0.1220 | 0.527 | 2 | |
| H-bonded to any TCR residue | 0.2174±0.4125 | 0.2574±0.1175 | 0.827 | 2 | |
| H-bonded to any non-CDR TCR residue | 0.0652±0.2238 | 0.1206±0.1151 | 0.664 | 2 | |
| In the edge [2] | 0.4783±0.4773 | 0.5588±0.0997 | 0.688 | 2 | |
| Closest distance to peptide (Å) | 3.7267±1.0947 | 5.1352±1.0994 | 5.15e-05 | 3 | *** |
| Number of H-bonds formed | 1.7200±1.2496 | 2.0264±0.7809 | 0.503 | 3 | |
| H-bonded to any CDR residue | 0.2800±0.4490 | 0.1714±0.0963 | 0.196 | 3 | |
| H-bonded to any peptide residue | 0.1200±0.3250 | 0.0620±0.1108 | 0.31 | 3 | |
| H-bonded to any TCR residue | 0.4000±0.4899 | 0.2491±0.1120 | 0.0866 | 3 | |
| H-bonded to any non-CDR TCR residue | 0.1200±0.3250 | 0.1197±0.1162 | 0.697 | 3 | |
| In the edge [2] | 0.2600±0.4271 | 0.5771±0.1191 | 0.00591 | 3 | *** |
| H-bonded to any peptide residue | 0.0495±0.1443 | 0.0659±0.1206 | 0.458 | all | |
| H-bonded to any CDR residue | 0.2050±0.3024 | 0.1682±0.0982 | 0.48 | all | |
| H-bonded to any TCR residue | 0.3401±0.3714 | 0.2372±0.1184 | 0.151 | all | |
| H-bonded to any non-CDR TCR residue | 0.1712±0.3112 | 0.1118±0.1283 | 0.355 | all | |
| In the edge [2] | 0.4459±0.4097 | 0.5874±0.1232 | 0.0795 | all | |
| Closest distance to peptide (Å) | 4.6398±1.7149 | 5.1926±1.2647 | 0.141 | all | |
| Number of H-bonds formed | 2.1126±1.4959 | 2.0031±0.9051 | 0.668 | all | |

1. Mean and standard deviation (for the 39 structures) of the proportion of residues that satisfy the property shown in the first column.

2. Three residues from the beginning and four from the end of the peptide. 3. In the last two properties, per-residue averages were used instead.

**Table S4.** TCR side attention analysis on all heads. In a cell, the left number is the mean value over the PDBIDs and the right number is the STD value.

| Property | Large Attention [1] | Small Attention [1] | p-value | Head | |
|---|---|---|---|---|---|
| H-bonded to any peptide residuetide | 0.0256 ± 0.1581 | 0.0853 ± 0.0577 | 0.043 | 0 | *** |
| H-bonded to any CDR residue | 0.6068 ± 0.4262 | 0.4135 ± 0.1061 | 0.0111 | 0 | *** |
| H-bonded to any TCR residue of own chain | 0.7051 ± 0.4038 | 0.6408 ± 0.0888 | 0.353 | 0 | |
| H-bonded to any CDR residue of own chain | 0.6068 ± 0.4262 | 0.3792 ± 0.0991 | 0.0027 | 0 | *** |
| H-bonded to any TCR residue of opposite chain | 0.1111 ± 0.2833 | 0.1565 ± 0.0756 | 0.371 | 0 | |
| H-bonded to any CDR residue of opposite chain | 0.0000 ± 0.0000 | 0.0642 ± 0.0673 | 8.37e-07 | 0 | *** |
| H-bonded to any TCR residue | 0.7308 ± 0.3897 | 0.7193 ± 0.0903 | 0.864 | 0 | |
| H-bonded to any non-CDR TCR residue | 0.1966 ± 0.3734 | 0.4479 ± 0.0738 | 0.000482 | 0 | *** |
| In the edge [2] | 0.8376 ± 0.3235 | 0.5904 ± 0.0387 | 5.1e-05 | 0 | *** |
| Closest distance to peptide (Å) | 9.1216 ± 3.1274 | 8.3528 ± 1.0097 | 0.119 | 0 | |
| Number of H-bonds formed | 1.7479 ± 1.1003 | 2.0919 ± 0.6806 | 0.0397 | 0 | *** |
| H-bonded to any peptide residuetide | 0.1453 ± 0.3162 | 0.0773 ± 0.0564 | 0.2 | 1 | |
| H-bonded to any CDR residue | 0.5171 ± 0.4333 | 0.4147 ± 0.1077 | 0.183 | 1 | |
| H-bonded to any TCR residue of own chain | 0.5427 ± 0.4315 | 0.6488 ± 0.0892 | 0.152 | 1 | |
| H-bonded to any CDR residue of own chain | 0.5085 ± 0.4335 | 0.3817 ± 0.0999 | 0.0913 | 1 | |
| H-bonded to any TCR residue of opposite chain | 0.1496 ± 0.3198 | 0.1510 ± 0.0714 | 0.979 | 1 | |
| H-bonded to any CDR residue of opposite chain | 0.0171 ± 0.1054 | 0.0616 ± 0.0678 | 0.0467 | 1 | *** |
| H-bonded to any TCR residue | 0.6368 ± 0.4480 | 0.7211 ± 0.0881 | 0.264 | 1 | |
| H-bonded to any non-CDR TCR residue | 0.1581 ± 0.2995 | 0.4511 ± 0.0648 | 5.28e-07 | 1 | *** |
| In the edge [2] | 0.5513 ± 0.4388 | 0.6055 ± 0.0443 | 0.477 | 1 | |
| Closest distance to peptide (Å) | 7.7849 ± 3.3952 | 8.4412 ± 0.9215 | 0.15 | 1 | |
| Number of H-bonds formed | 1.9957 ± 1.5568 | 2.0725 ± 0.6687 | 0.742 | 1 | |
| H-bonded to any peptide residuetide | 0.1127 ± 0.2935 | 0.0805 ± 0.0585 | 0.654 | 2 | |
| H-bonded to any CDR residue | 0.3775 ± 0.4470 | 0.4247 ± 0.0992 | 0.614 | 2 | |
| H-bonded to any TCR residue of own chain | 0.4853 ± 0.4615 | 0.6499 ± 0.0851 | 0.067 | 2 | |
| H-bonded to any CDR residue of own chain | 0.3480 ± 0.4379 | 0.3919 ± 0.0947 | 0.633 | 2 | |
| H-bonded to any TCR residue of opposite chain | 0.1324 ± 0.3278 | 0.1557 ± 0.0707 | 0.759 | 2 | |
| H-bonded to any CDR residue of opposite chain | 0.0588 ± 0.2353 | 0.0614 ± 0.0678 | 0.933 | 2 | |
| H-bonded to any TCR residue | 0.5882 ± 0.4451 | 0.7249 ± 0.0874 | 0.127 | 2 | |
| H-bonded to any non-CDR TCR residue | 0.3235 ± 0.4021 | 0.4399 ± 0.0748 | 0.144 | 2 | |
| In the edge [2] | 0.4706 ± 0.4705 | 0.6060 ± 0.0390 | 0.126 | 2 | |
| Closest distance to peptide (Å) | 7.9293 ± 4.0178 | 8.3982 ± 1.0153 | 0.588 | 2 | |
| Number of H-bonds formed | 2.1765 ± 1.8190 | 2.0746 ± 0.6642 | 0.607 | 2 | |
| H-bonded to any peptide residuetide | 0.0877 ± 0.2470 | 0.0818 ± 0.0572 | 0.909 | 3 | |
| H-bonded to any CDR residue | 0.4342 ± 0.4318 | 0.4234 ± 0.1013 | 0.852 | 3 | |
| H-bonded to any TCR residue of own chain | 0.5921 ± 0.4270 | 0.6499 ± 0.0865 | 0.466 | 3 | |
| H-bonded to any CDR residue of own chain | 0.3947 ± 0.4161 | 0.3908 ± 0.0969 | 0.927 | 3 | |
| H-bonded to any TCR residue of opposite chain | 0.2237 ± 0.2993 | 0.1476 ± 0.0693 | 0.116 | 3 | |
| H-bonded to any CDR residue of opposite chain | 0.0526 ± 0.1916 | 0.0615 ± 0.0631 | 0.767 | 3 | |
| H-bonded to any TCR residue | 0.6842 ± 0.3723 | 0.7236 ± 0.0884 | 0.57 | 3 | |
| H-bonded to any non-CDR TCR residue | 0.3684 ± 0.3590 | 0.4391 ± 0.0678 | 0.263 | 3 | |
| In the edge [2] | 0.7281 ± 0.3735 | 0.5962 ± 0.0430 | 0.0464 | 3 | *** |
| Closest distance to peptide (Å) [3] | 8.6919 ± 3.5172 | 8.4100 ± 1.0378 | 0.597 | 3 | |
| Number of H-bonds formed [3] | 2.1535 ± 1.4009 | 2.0690 ± 0.6701 | 0.622 | 3 | |
| H-bonded to any peptide residue | 0.0862±0.1368 | 0.0805±0.0675 | 0.828 | all | |
| H-bonded to any CDR residue | 0.4846±0.2216 | 0.4103±0.1040 | 0.0478 | all | *** |
| H-bonded to any TCR residue of own chain | 0.6013±0.1999 | 0.6561±0.0880 | 0.117 | all | |
| H-bonded to any CDR residue of own chain | 0.4643±0.2180 | 0.3752±0.0922 | 0.0107 | all | *** |
| H-bonded to any TCR residue of opposite chain | 0.1679±0.1714 | 0.1497±0.0793 | 0.562 | all | |
| H-bonded to any CDR residue of opposite chain | 0.0306±0.0857 | 0.0672±0.0743 | 0.0369 | all | *** |
| H-bonded to any TCR residue | 0.6845±0.1650 | 0.7294±0.0880 | 0.0987 | all | |
| H-bonded to any non-CDR TCR residue | 0.2940±0.1923 | 0.4672±0.0846 | 3.88e-05 | all | *** |
| In the edge [2] | 0.6434±0.2064 | 0.5928±0.0570 | 0.218 | all | |
| Closest distance to peptide (Å) [3] | 8.4072±2.2892 | 8.4122±0.9592 | 0.988 | all | |
| Number of H-bonds formed [3] | 2.0234±0.9370 | 2.0875±0.6685 | 0.589 | all | |

1. Mean and standard deviation (for the 39 structures) of the proportion of residues that satisfy the property shown in the first column.
2. Four residues from the beginning and four from the end of the CDR. 3. In the last two properties, per-residue averages were used instead.

### PYMOL COMMAND FOR PDB 5TEZ

```
fetch 5TEZ;
set seq_view, 1;
bg_color white;
hide all;
remove waters;
select beta, chain J and not solvent;
select alpha, chain I and not solvent;
select mhc, (chain A or chain B or chain D or chain E) and not
   solvent;
show cartoon, alpha;
color wheat, alpha;
show cartoon, beta;
color lightblue, beta;
show cartoon,mhc
color grey90, mhc;
create obj_mhc, mhc
show surface, obj_mhc
set transparency=0.2
sel beta_cdr3, (chain J and resi 91:104);
#set cartoon_side_chain_helper, on
#show sticks, beta_cdr3;
#util.cbag beta_cdr3;
color palecyan, beta_cdr3
sel alpha_cdr3, (chain I and resi 91:105);
#set cartoon_side_chain_helper, on
#show sticks, alpha_cdr3;
#util.cbag alpha_cdr3;
color lightpink, alpha_cdr3
select epitope, chain C and not solvent;
show sticks, epitope;
color yellow, epitope
#util.cbay epitope;
#select cdr3, alpha_cdr3 or beta_cdr3;
#select tcr, alpha or beta;
#dist H_cdr_p, cdr3, epitope, mode=2;
#hide labels, H_cdr_p;
#color black, H_cdr_p;
#dist H_cdr_tcr, cdr3, tcr, mode=2;
#hide labels, H_cdr_tcr;
#color grey, H_cdr_tcr;
sel atten_a_head1, (resi 104 and chain I);
#color pink, atten_a_head1;
sel atten_a_head2, (resi 101 and chain I);
#color pink, atten_a_head2;
```

```
sel atten_a_head3, (resi 101 and chain I);
#color pink, atten_a_head3;
sel atten_b_head0, (resi 99 and chain J);
#color pink, atten_b_head0;
show sticks, atten_a_head1
show sticks, atten_a_head2
show sticks, atten_a_head3
show sticks, atten_b_head0
color magenta, atten_a_head1
color magenta, atten_a_head2
color magenta, atten_a_head3
color cyan, atten_b_head0
sel atten_1_int, (resi 93 and chain I)
sel atten_23_int, (resi 94 and chain I)
sel atten_230_int, (resi 6 and chain C)
show sticks, atten_1_int
show sticks, atten_23_int
show sticks, atten_0_int
sel int_int,(resi 98 and chain J)
show sticks, int_int
color atomic,(not elem C)
color gray90, obj_mhc
dist a1_hb,(resi 104 and chain I),(resi 93 and chain I),mode=2
dist a23_hb,(resi 101 and chain I),(resi 94 and chain I),mode=2
dist a23p_hb,(resi 101 and chain I),(resi 6 and chain C),mode=2
dist a0_hb,(resi 99 and chain J),(resi 6 and chain C),mode=2
dist intint,(resi 94 and chain I),(resi 98 and chain J),mode=2
hide labels, a1_hb
hide labels, a23_hb
hide labels, a23p_hb
hide labels, a0_hb
hide labels, intint
```
